# Intestinal receptor of SARS-CoV-2 in inflamed IBD tissue is downregulated by HNF4A in ileum and upregulated by interferon regulating factors in colon

**DOI:** 10.1101/2020.06.24.169383

**Authors:** Bram Verstockt, Sare Verstockt, Saeed Abdu Rahiman, Bo-jun Ke, Kaline Arnauts, Isabelle Cleynen, João Sabino, Marc Ferrante, Gianluca Matteoli, Séverine Vermeire

**Affiliations:** Department of Gastroenterology and Hepatology, University Hospitals Leuven, KU Leuven, Leuven, Belgium; Department of Chronic Diseases, Metabolism and Ageing (CHROMETA), Translational Research Center for Gastrointestinal Disorders (TARGID), KU Leuven, Leuven, Belgium; Department of Development and Regeneration, Stem Cell Institute Leuven (SCIL), KU Leuven, Leuven, Belgium; Department of Human Genetics, Laboratory for Complex Genetics, KU Leuven, Leuven, Belgium

**Author notes:** contributed equally (Bram Verstockt, Sare Verstockt). contributed equally (Gianluca Matteoli, Séverine Vermeire). **Corresponding author**, Séverine Vermeire, MD, PhD, Department of Gastroenterology and Hepatology, University Hospitals Leuven, Herestraat 49 3000 Leuven, Belgium. @ibdleuven @TARGID_KULEUVEN @bverstockt @SareVerstockt @gmatteoli1978 @JoaoPGSabino @ICleynen @saeedfc.

**Keywords:** COVID-19, ACE2, TMPRSS2, inflammatory bowel diseases, SARS-CoV-2, HNF4A, interferon, organoids, transcriptomics, single cell, intestinal inflammation

## Abstract

Patients with IBD are considered immunosuppressed, but do not seem more vulnerable for COVID-19. Nevertheless, intestinal inflammation has shown an important risk factor for SARS-CoV-2 infection and prognosis. Therefore, we investigated the effect of intestinal inflammation on the viral intestinal entry mechanisms, including *ACE2*, in IBD.

We collected (un)inflamed mucosal biopsies from CD (n=193) and UC (n=158) patients, and 51 matched non-IBD controls for RNA sequencing, differential gene expression and co-expression analysis. Organoids from UC patients were subjected to an inflammatory mix and processed for RNA sequencing. Transmural ileal biopsies were processed for single-cell (sc) sequencing. Publicly available colonic sc-RNA sequencing data, and microarrays from tissue pre/post anti-TNF therapy, were analyzed.

In inflamed CD ileum, *ACE2* was significantly decreased compared to control ileum (p=4.6E-07), whereas colonic *ACE2* expression was higher in inflamed colon of CD/UC compared to control (p=8.3E-03; p=1.9E-03). Sc-RNA sequencing confirmed this *ACE2* dysregulation, and exclusive epithelial *ACE2* expression. Network analyses highlighted *HNF4A* as key regulator of ileal *ACE2*, while pro-inflammatory cytokines and interferon regulating factors regulated colonic *ACE2.* Inflammatory stimuli upregulated *ACE2* in UC organoids (p=1.7E-02), not in non-IBD controls (p=9.1E-01). Anti-TNF therapy restored colonic *ACE2* dysregulation in responders.

Intestinal inflammation alters SARS-CoV-2 coreceptors in the intestine, with opposing effects in ileum and colon. *HNF4A*, an IBD susceptibility gene, is an important upstream regulator of *ACE2* in ileum, whereas interferon signaling dominates in colon. Our data support the importance of adequate control of IBD in order to reduce risk of (complicated) COVID-19.

## INTRODUCTION

Since the novel betacoronavirus SARS-CoV-2 was first reported in the province of Wuhan, China, at the end of 2019, the virus has spread over more than 200 countries, causing more than 8.9 million infections, including almost 470.000 death globally.^1^ Despite being primarily a respiratory virus, coronavirus disease 2019 (COVID-19) can also present with non-respiratory signs, including digestive symptoms as diarrhea, nausea and ageusia.^2, 3, 4^

Although it is thought that SARS-CoV-2 primarily infects the lungs with transmission via the respiratory route, the gastro-intestinal tract can be an alternative viral target organ. Indeed, the SARS-CoV-2 receptor angiotensin converting enzyme 2 (ACE2) is highly expressed on differentiated enterocytes, with strong induction of generic viral response programs upon viral binding.^5, 6^ The cellular entry of coronaviruses depends on the binding of the spike (S) protein to a specific receptor, followed by an S protein priming by proteases, with key players ACE2 (receptor for the S protein) and TMPRSS2 (protease) in case of COVID-19.^6, 7, 8^ Furthermore, based on protein crystal structures, data predicted that the Middle East respiratory syndrome (MERS)-CoV receptor dipeptidyl peptidase 4 (DDP4) might act as a candidate binding target or co-receptor of SARS-CoV-2.^9, 10^ In line, proteomic studies in COVID-19 patients suggested a prognostic role for DDP4.^11^ Upon cellular entry in nasal goblet secretory cells, lung type II pneumocytes and ileal absorptive enterocytes, an interferon-driven mechanism is initiated, including the upregulation of *ACE2* which further enhances infection.^7^

Why *ACE2*, the S protein receptor, is abundantly expressed on intestinal epithelium, is not entirely understood. Recent studies have addressed the homeostatic role of ACE2 on intestinal epithelial cells demonstrating defective intestinal amino acid absorption in ACE2 deficient mice.^12^ Mechanistically, ACE2 independently of its role on the renin angiotensin system (RAS), is essential for regulating epithelial tryptophan absorption, expression of antimicrobial peptides, and consequently regulating the ecology of the gut microbiome promoting homeostasis and preventing intestinal inflammation.^13^ Thus, ACE2 regulation could be link to the pathogenesis of IBD, playing a role as modulator of epithelial immune homeostatic functions.

Individual susceptibility to COVID-19 may correlate with the expression of these designated (co)receptors. In this respect, it is unknown how inflammation affects *ACE2, TMPRSS2* and/or *DDP4* expression in ileum and colon. Inflammatory Bowel Disease (IBD) is a prototype gastrointestinal disease characterized by a chronic relapsing inflammatory infiltrate of the small and/or large bowel. So far, data on COVID-19 in patients with IBD are rather limited,^14, 15, 16, 17, 18^ although suggest that increasing age, a diagnosis of ulcerative colitis [UC] (as opposed to Crohn’s disease [CD]) and increasing disease activity are linked with a more severe course of COVID-19. In contrast, anti-inflammatory IBD therapy has not yet been associated with COVID-19 risk. Using a combination of bulk and single cell transcriptomics and organoid cultures, we studied the intestinal expression of several SARS-CoV-2 co-receptors in the healthy gut and in IBD and investigated whether inflammation alters co-receptor expression.

## MATERIALS AND METHODS

### Patients

This study was carried out at the University Hospitals Leuven (Leuven, Belgium). All included patients had given written consent to participate in the Institutional Review Board approved IBD Biobank of University Hospitals Leuven, Belgium (B322201213950/S53684 and B322201110724/S52544). Endoscopy-derived (un)inflamed mucosal biopsies were obtained cross-sectionally from IBD patients requiring colonoscopy during routine care (**Supplementary Table S1**). Samples from individuals undergoing colonoscopy for polyp detection were included as controls. Transmural ileal biopsies, derived during right hemicolectomy from CD patients and patients with colorectal cancer (CRC), were collected, stored in RPMI-1640 medium on ice until single cell isolation.

### Organoids

Mucosal biopsies from both uninflamed and macroscopically inflamed colon segments (UC only) were processed as reported earlier.^19^ In brief, crypts were isolated and cultured as organoids for at least four weeks. Inflammation was then re-induced using an inflammatory mix (100 ng/ml TNF-α, 20 ng/ml IL-1β, 1μg/ml Flagellin) during 24 hours.^19^

### Bulk transcriptomics

Inflamed biopsies were taken at the most affected site at the edge of an ulcerative surface, whereas uninflamed biopsies were taken randomly in macroscopic unaffected areas. All were stored in RNALater buffer (Ambion, Austin, TX, USA) and preserved at −80°C. As described previously,^20^ RNA from biopsies was isolated using the AllPrep DNA/RNA Mini kit (Qiagen, Hilden, Germany), and RNA libraries were prepared using the TruSeq Stranded mRNA protocol (Illumina, San Diego, USA). RNA from organoids was extracted using the RNeasy Mini Kit (Qiagen) and libraries were constructed by the Lexogen QuantSeq 3’ mRNA-Seq Library Kit FWD (Lexogen, Vienna, Austria).^19^ All RNA libraries were sequenced by the Illumina HiSeq4000 (Illumina, San Diego, CA). Raw sequencing data were aligned to the reference genome (GRCh37) using Hisat2 (version 2.1.0) ^21^ and absolute counts generated using HTSeq.^22^ Counts were normalized and protein coding genes selected (Ensemble hg 19 reference build)^23^ using the DESeq2 package.^24^ A weighted gene co-expression network (WGCNA) was generated,^25^ as described earlier.^26, 27^ The module eigengene was defined as the first principal component summarizing the expression patterns of all genes into a single expression profile within a given module. Genes showing the highest correlation with the module eigengene were referred to as hub genes. Pathway and upstream regulator analyses were performed using Ingenuity Pathway Analysis (IPA, QIAGEN, Aarhus, Denmark), with network visualization via Cytoscape (v3.8.0).^28^ Publicly available microarray datasets of ileal and colonic biopsies (GEO GSE14580, GSE12251, GSE16879) were accessed to investigate the effect of anti-TNF therapy on genes of interest.^29, 30^

### Single cell transcriptomics

Transmural ileal samples were treated with 1mM DTT and 1mM EDTA in 1x Hank’s balanced salt solution (HBSS), and 1 mM EDTA in HBSS at 37°C for 30 minutes, respectively. Then, tissue was transferred into a sterile gentleMACS C tube (Miltenyi Biotec), and digested with 5.4 U/mL collagenase D (Roche applied science), 100 U/mL DNase I (Sigma) and 39.6 U/mL dispase II (Gibco) with the gentleMACS™ Dissociator (program human_tumor_02.01). Samples were incubated for 30 minutes at 37°C at 250 rpm. Dissociated samples were filtered with 70 μm cell strainers, and treated with red blood cell lysis buffer (11814389001, Roche). After centrifugation, single-cell suspensions were re-suspended in 0.4% BSA in PBS, and were immediately processed with 10x 3’ v3 GEM kit, and loaded on a 10x chromium controller to create Single Cell Gel beads in Emulsion (GEM). A cDNA library was created and assessed using a 10x 3’ v3 library kit, and was then sequenced on a NovaSeq 6000 system (Illumina). Pre-processing of the samples including alignment, and counting was performed using Cell Ranger Software from 10x (Version: 3.0.2).

Publicly available colonic single cell RNA sequencing data (sc-RNA seq) (Single Cell Portal, SCP 259) were downloaded and visualized using the SCP data browser.^31^ For colonic epithelial single cell data, tSNE coordinates and publicly available annotation with the data was used for visualization and analysis.

Annotation of the ileal data was performed using SingleR R package, with inbuilt Human Cell Atlas data as reference. Quality control, clustering and dimensionality reduction of sc-RNA seq data was performed using Seurat R package (Version 3.1.5).^32, 33^ Data from each 10x run were integrated after performing SCTransform on each dataset, with percentage of mitochondrial genes set as a parameter to be regressed. Single Cell Network Inference (SCENIC) analysis was performed using a python implementation of the SCENIC pipeline (PySCENIC) (Version 0.9.19).^34^

### Immunofluorescence staining

Transmural ileal biopsies, obtained during abdominal surgery in patients with IBD and CRC, were fixed in 4% formalin, embedded in paraffin and sections of 5μm were cut (Translational Cell & Tissue Research Laboratory, University Hospitals Leuven and at VIB & KU Leuven Center for Brain & Disease Research). After deparaffinization, antigen retrieval was done in Tris-EDTA buffer (10 mM Tris base, 1 mM EDTA solution, 0.05% Tween 20, pH 9.0) at 95°C for 30 minutes. 1% BSA in PBST (0.1% tween-20 and 0.5% sodium azide) was used to block non-specific binding of detection antibodies and gently permeabilize before ACE2 and Cytokeratin AE1/AE3 staining. In brief, ACE2 (Polyclonal, Cell Signaling Technology) and cytokeratin (IgG1-kappa, clone AE1/AE3, Dako) were applied in 1% BSA, followed by donkey anti-rabbit Cy3 (Jackson Immuno Research) and donkey anti-mouse Alexa fluor 488 (Invitrogen). Slides were mounted in SlowFade™ Diamond Antifade Mountant (Invitrogen), and stored at 4 °C before imaging. Images were acquired using a Zeiss LSM 780 at the Cell and Tissue Imaging Cluster (CIC) at KU Leuven.

### Genetics

All samples were genotyped using the Illumina GSA array. All SNPs and samples with more than 10% missingness rate were removed, as were SNPs with minor allele frequency (MAF)<0.001. Genotypes for rs6017342 *(HNF4A)* were extracted. All steps were performed using PLINK (v1.90b4.9).^35^

### Statistical analysis

Statistical analysis was performed using R 3.6.2 (The R foundation, Vienna, Austria). Pearson correlation coefficients were computed to assess the correlation between individual genes. Multivariate analysis was performed using the R package “lm.beta”. Continuous variables on graphs were expressed as median and interquartile range (IQR). *ACE2, DPP4* and *TMPRSS2 c*omparisons were done using two sample t-tests or Wilcoxon tests, as appropriate. In case of hypothesis-free comparisons, ie. genome-wide differential gene expression analyses, multiple testing correction was applied (adjusted p [adj. p], Benjamini-Hochberg method).

## RESULTS

### Intestinal *ACE2, TMPRSS2* and *DPP4* expression in IBD patients versus non-IBD controls

First, we studied the expression patterns of *ACE2, DPP4* and *TMPRSS2* in ileum and colon biopsies from 351 IBD patients (193 CD, 158 UC) and 51 non-IBD controls based on bulk RNA sequencing.

In non-IBD controls, *ACE2* and *DPP4* expression levels were strongly increased in ileum compared to colon (fold change (FC) =32.0, p=6.3E-13; FC=16.5, p=6.3E-13) (**Figure 1A-B**). In contrast, ileal *TMPRSS2* was lower compared to colon (FC=-2.9, p=6.3E-13) (**Figure 1C**).

**Figure 1:**
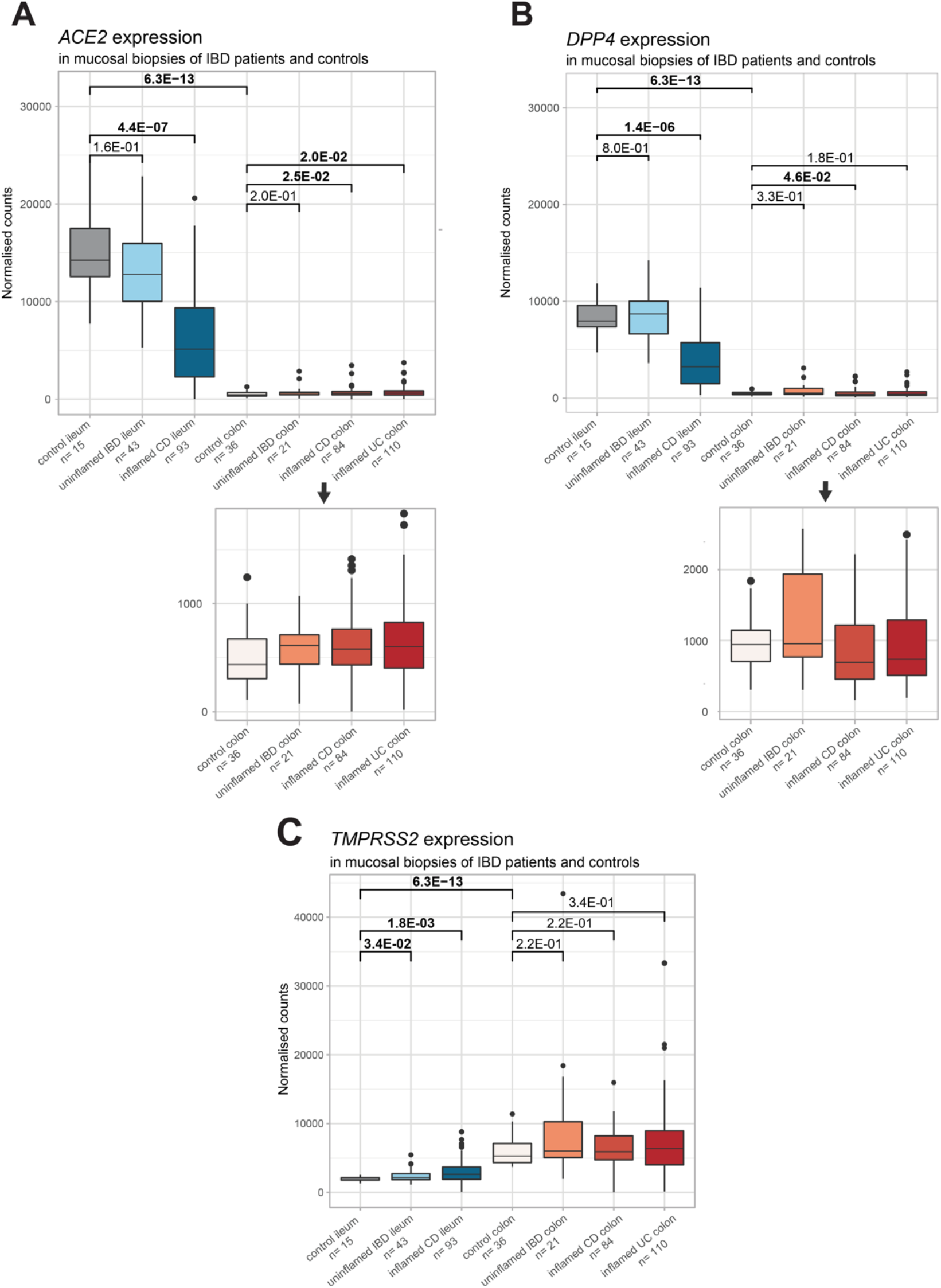
Mucosal *ACE2, DPP4* and *TMPRSS2* in IBD patients and controls. (**A**) Boxplots of mucosal *ACE2* as measured by RNA sequencing (normalized counts). (**B**) Boxplots of mucosal *DPP4* as measured by RNA sequencing (normalized counts). (**C**) Boxplots of mucosal *TMPRSS2* as measured by RNA sequencing (normalized counts). Significant comparisons are highlighted in bold. CD, Crohn’s disease; control, non-IBD controls; IBD, inflammatory bowel disease; UC, ulcerative colitis

When turning to tissue from IBD patients, *ACE2* and *DPP4* levels in uninflamed IBD ileum were similar to those observed in matched control ileum (p=1.6E-01; p=8.0E-01) (**Figure 1A-B**). *TMPRSS2* however, was significantly upregulated compared to control ileum (FC=1.2, p=3.4E-02) (**Figure 1C**). In uninflamed IBD colon, expression levels of *ACE2, DDP4* and *TMPRSS2* did not differ from control colon (p=2.0E-01; p=3.3E-01; 2.2E-01) (**Figure 1A-C**).

In inflamed CD ileum, *ACE2* and *DPP4* expression was significantly decreased compared to control ileum (FC=-2.8, p=4.4E-07; FC=-2.5, p=1.4E-06) (**Figure 1A-B**). *TMPRSS2* behaved opposite, with a significant upregulation in inflamed ileum versus control ileum (FC=1.4, p=1.8E-03) (**Figure 1C**). At colonic level, *ACE2* expression was higher in inflamed CD and UC colon than in control colon (FC=1.4, p=2.5E-02; FC=1.4, p=2.0E-02 respectively) (**Figure 1A**). Except for a decrease in *DPP4* expression in inflamed CD colon versus control colon (FC=1.3, p=4.6E-02), no dysregulations were observed for colonic *DPP4* and *TMPRSS2* (p=1.8E-01; p≤3.4E-01) (**Figure 1B-C**).

Despite *ACE2* being X-linked, multivariate analysis did not reveal any contribution of sex to mucosal *ACE2* expression (p=5.1E-01), nor of age (p=1.4E-01), diagnosis (p=5.6E-01) or disease duration (p=5.2E-01). Intestinal *ACE2* expression was significantly affected by biopsy location (p=2.8E-34) and inflammatory state (p=4.2E-12).

### Gene co-expression analysis of the *ACE2-, DPP4-* and *TMPRSS2-related* networks

To get a better understanding of the biological network of *ACE2*, *DPP4* and *TMPRSS2*, we performed WGCNA on all mucosal biopsies.

At ileal level, we identified 18 co-expression modules (clusters) ranging in size from 106 to 1465 genes (**Figure 2A)**. One module contained both *ACE2* and *DPP4* (module “blue”; 1134 genes) **(Supplementary Table S2)**. The strongest correlation with the eigengene (i.e. the principal component) of this *ACE2/DPP4*-module was found for hub genes *MMP5* (r=0.94, p=8.6E-74), *ZNF664* (r=0.94, p=3.7E-71) and *DPP4* (r=0.93, p=1.2E-68) (**Figure 2A**). Moreover, *ACE2* also seemed to have a central role in this co-expression network with a correlation value of r=0.86 (p=4.6E-45) (**Figure 2A**). Pathway analysis of the *ACE2/DPP4-* module found enrichment for epithelium-related metabolic pathways such as Xenobiotic Metabolism Signaling, Nicotine Degradation II and Melatonin Degradation (p<1.0E-08). Predicted upstream analysis (using curated datasets in IPA) highlighted the transcription regulator HNF4A, an IBD susceptibility gene, as the most likely upstream regulator of the *ACE2/DPP4-module* (p=1.2E-11).

**Figure 2:**
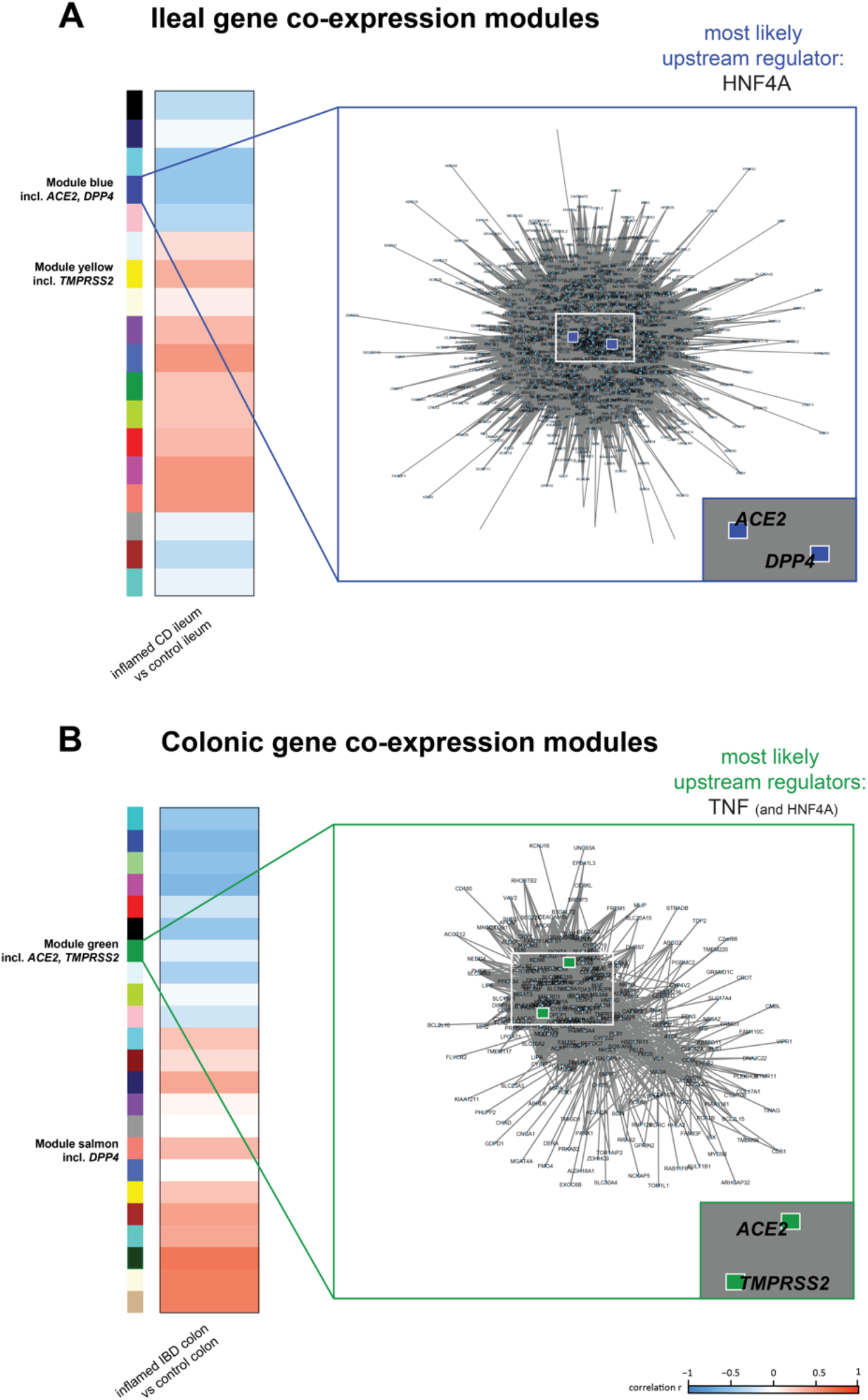
Gene co-expression analysis of the ACE2-, DPP4- and TMPRSS2-related networks. WGCNA was applied on mucosal biopsies of (**A**) IBD ileum and non-IBD control ileum, and (**B**) IBD colon and non-IBD control colon. A positive correlation (highlighted in red) means an upregulation in disease as compared to matched controls, while a negative correlation (highlighted in blue) refers to a downregulation in disease. The reported networks visualize genes having a correlation value of ≥ 0.85 with the module eigengene. CD, Crohn’s disease; control, non-IBD controls; IBD, inflammatory bowel disease; incl., including; WGCNA, Weighted Gene Co-expression Network Analysis;

*TMPRSS2* belonged to a separate module “yellow” (1126 genes) with hub gene *COA3* (r=0.92, p=4.7E-61) (**Figure 2A, Supplementary Table S3**). Genes within this module were mainly related to mitochondrial functions (eg. Oxidative Phosphorylation, Mitochondrial dysfunction and Sirtuin Signaling, p<1.6E-29), and their top upstream regulator was again HNF4A (p=1.5E-27).

At colonic level, 24 co-expression modules were present ranging in size from 128 to 2267 genes (**Figure 2B**). In contrast to the ileum, colonic *ACE2* and *DPP4* were not co-expressed (**Supplementary Tables S3**), with *ACE2* being part of module “green” (797 genes). Here, *ACE2* co-clustered with *TMPRSS2.* The *ACE2*-module with top hub gene *TMEM63B* (r=0.89, p=5.8E-81) did not show significant enrichment for specific pathways. Upstream analysis of this module ranked TNF and again HNF4A as the top regulators (p=7.7E-06; p=9.4E-03).

Lastly, we studied the relationship between mucosal *ACE2* and *HNF4A* expression levels. Ileal *ACE2* expression strongly correlated with ileal *HNF4A* expression (r=0.69, p<2.2E-16), whereas colonic levels showed limited correlation (r=0.2, p=1.3E-03) (**Supplementary Figure S1**).

### Single nucleotide polymorphisms in *HNF4A* linked to *ACE2* expression in ileum but not in colon

As the expression of *ACE2-*modules was found to be driven by the IBD susceptibility locus, *HNF4A*, we next studied the genetic variability in rs6017342 (i.e. the causal IBD variant in this locus, ^36^), and its relationship with *ACE2* and *HNF4A* expression, both in inflamed ileum and colon. Ileal *ACE2* levels were lower in patients carrying the *HNF4A-AA* genotype, compared to patients carrying the C-allele, i.e. *HNF4A-* AC or *HNF4A-CC* genotypes (p=2.8E-02) (**Figure 3**). Colonic *ACE2* expression was independent of the *HNF4A* genotype (p=6.7E-01).

**Figure 3:**
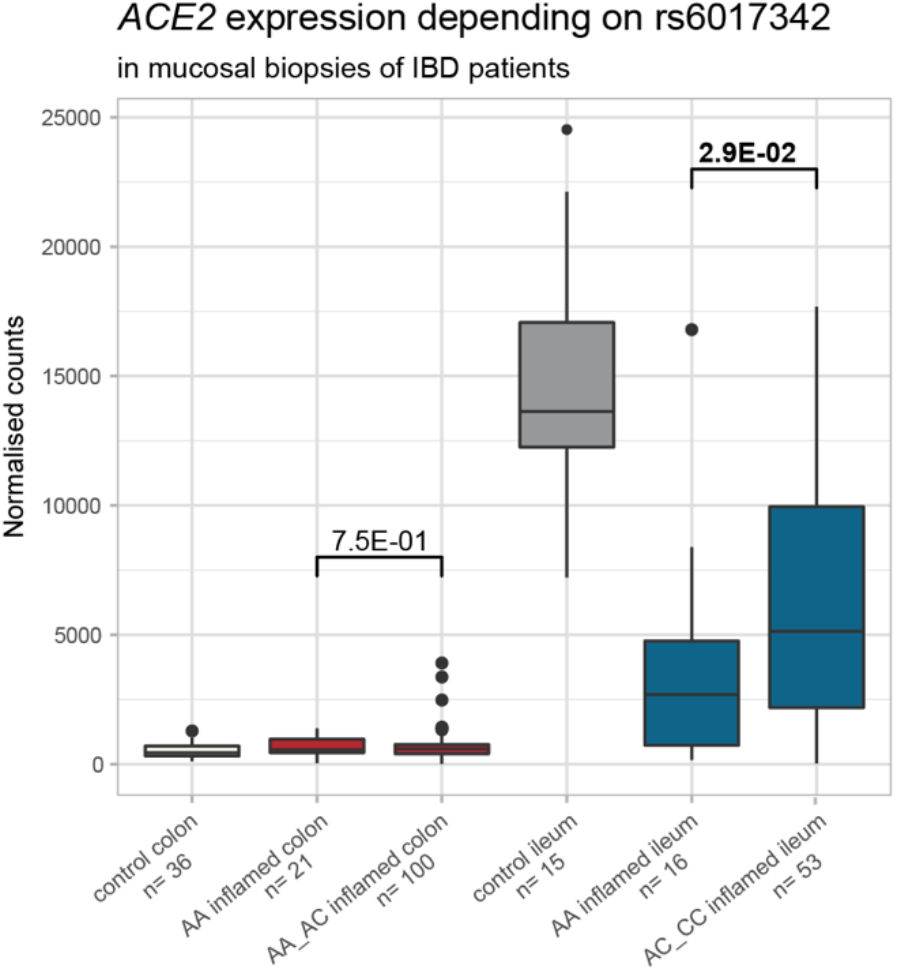
Mucosal *ACE2* and *HNF4A* in IBD patients and controls depending on rs6017342 genotype. Boxplots of mucosal *ACE2* as measured by RNA sequencing (normalized counts). Significant comparisons are highlighted in bold. controls, non-IBD controls

### Decrease of ACE2/TMPRSS2 double positive cells in inflamed ileum, but not in colon

*ACE2* expression in the gastrointestinal tract is primarily found in absorptive enterocytes,^7, 37^ which could indirectly be confirmed through the significant correlation (p<2.2E-16) between mucosal *ACE2* and several epithelial marker genes *(APOA1, SI, FABP6, ENPEP)* (**Supplementary Figure S2**). To further examine the expression of genes associated with risk of SARS-CoV-2 infection in IBD patients, we employed sc-RNA seq to profile transmural biopsies of (un)inflamed regions of resected tissue from six CD patients undergoing ileocaecal resection. Unaffected ileal tissue from five patients with CRC undergoing right hemicolectomy was used as control. A total of 78,722 cells were used for downstream analyses containing a similar number of cells from each type of tissue (inflamed CD, uninflamed CD and healthy tissue) (**Supplementary Figure S3B**). Sixty-one cell clusters belonging to epithelial, immune and stromal cells were obtained using unsupervised clustering (**Figure 4A, Supplementary Figure S3A**). Cell clusters were annotated by correlating the cluster gene expression profiles with Human Cell Atlas using SingleR, as previously described.^38^ *ACE2* expression was found exclusively in epithelial cell clusters (**Figure 4B-C**), which could also be confirmed using immunofluorescence staining (**Figure 5**). To define the epithelial cell subtypes expressing *ACE2* at deeper resolution, clusters annotated as epithelial cells by SingleR were extracted and re-clustered (**Figure 4D**). The re-clustered epithelial cell subtypes were annotated using a marker panel designed based on previous reports (**Supplementary Figure S3C**).^39^ Three enterocyte clusters were identified, out of which two clusters co-expressed *ACE2, TMPRSS2* and *DPP4.* Most prominent *ACE2* expression was observed in the ACE2/TMPRSS2 Enterocytes 1 cluster (**Figure 4G-I, Supplementary Figure S3D**).

**Figure 4:**
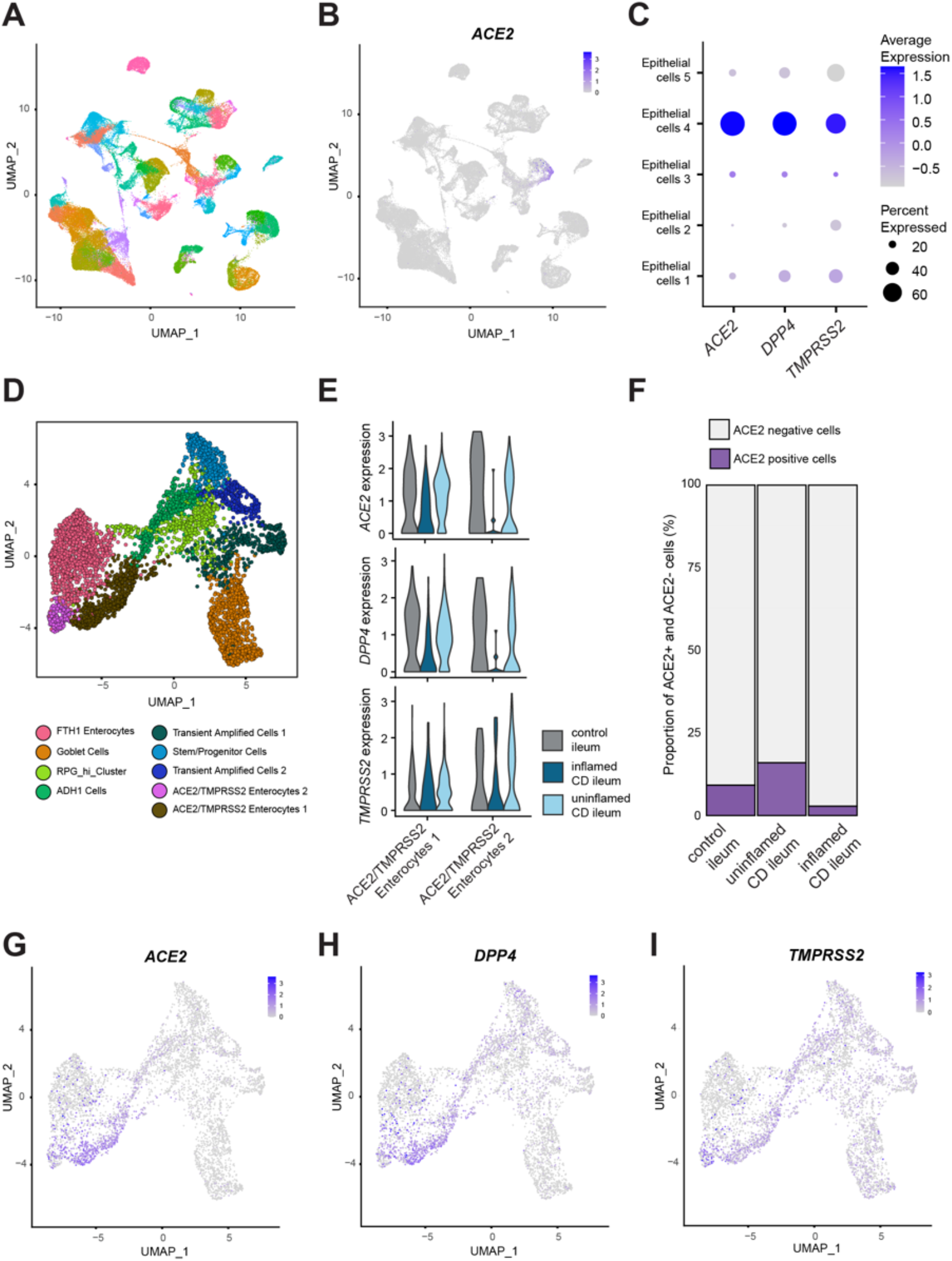
Decrease of ACE2/TMPRSS2 double positive cells in inflamed ileum in CD patients. (**A**) Uniform Manifold Approximation and Projection (UMAP) plot showing unsupervised clustering of integrated single cell RNA sequencing data from control, uninflamed and inflamed ileal tissue. (**B**) Expression of *ACE2* overlaid on the UMAP plot as in A. (**C**) Expression of *ACE2, DPP4*, and *TMPRSS2* in ileal epithelial cells. (**D**) UMAP showing epithelial sub clusters obtained upon re-clustering only the epithelial cells in ileum. (**E**) Expression and distribution of *ACE2, DPP4*, and *TMPRSS2* in the two enterocyte clusters co expressing *ACE2* and *TMPRSS2* split between control, uninflamed and inflamed samples. (**F**) Proportion of ACE2+ and ACE2-cells in control, uninflamed and inflamed samples in the ileal epithelial cells. (**G-I**) Gene expression overlaid on the UMAP Plot as in panel D of *ACE2*, *DPP4* and *TMPRSS2* respectively. CD, Crohn’s disease; control, non-IBD controls

Next, we asked whether *ACE2* expression varied across ileal tissue with inflammatory state, as observed in our bulk transcriptomic data (**Figure 1A**). *ACE2* expression and frequency of ACE2 positive cells were clearly reduced in ileum of patients with active CD, compared to uninflamed or healthy tissue (**Figure 4E-F, Supplementary Figure S3E**). A similar reduction of *DPP4* expression was observed in the inflamed samples in the ACE2/TMPRSS2 Enterocytes 1 and ACE2/TMPRSS2 Enterocytes 2 clusters (**Figure 4E**). In line, reduction of ACE2 expression in inflamed ileum compared to healthy tissue was also confirmed with confocal imaging (**Figure 5**).

**Figure 5:**
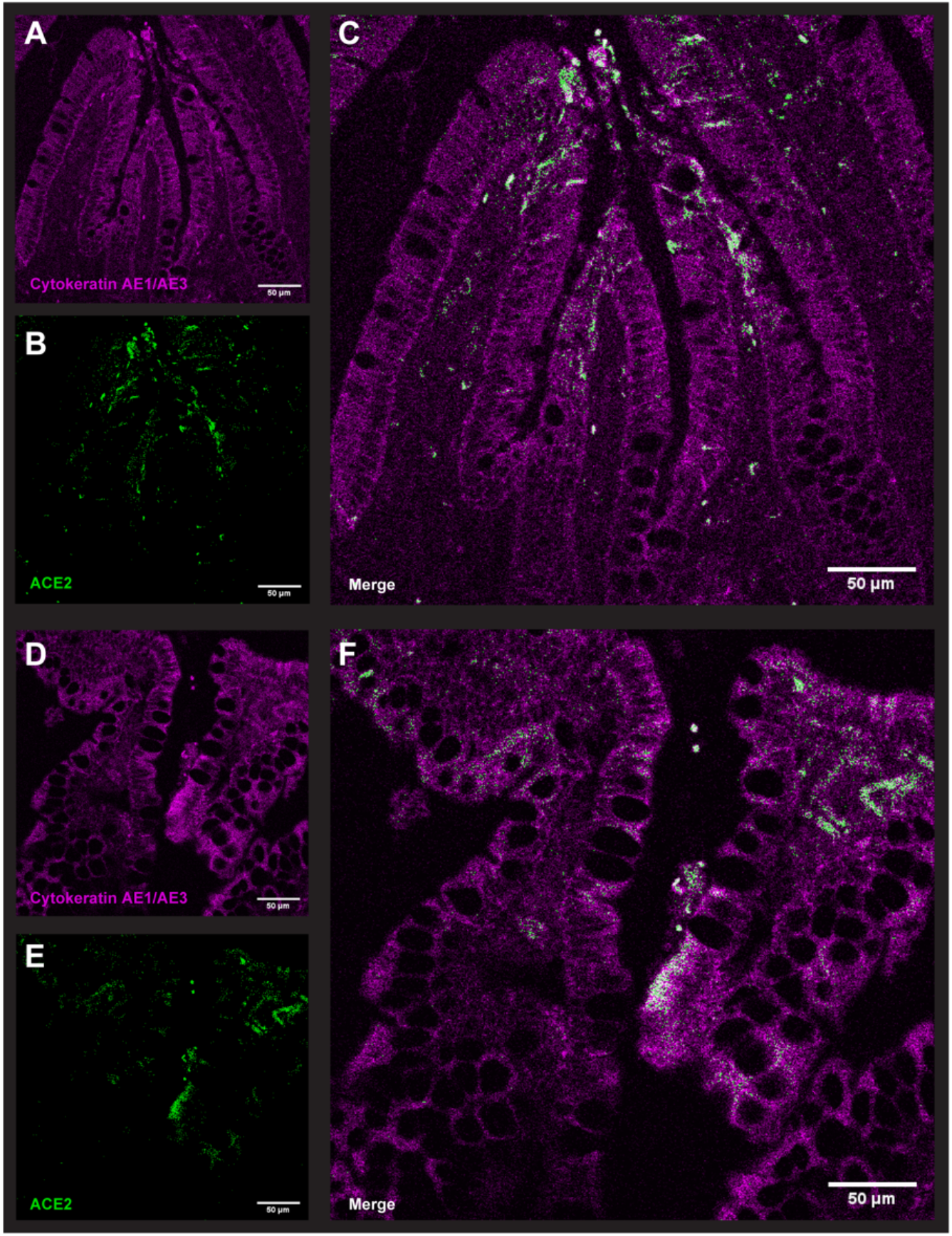
Cytokeratin AE1/AE3 and ACE2 expression in human gut. Confocal microscopy images of human gut in which ACE2 positive epithelial cells were stained with cytokeratin AE1/AE3 (magenta) and ACE2 (green). The scale bar in the immunofluorescent image represents 50 μm. Normal ileum from patient with colorectal cancer (**A-C**); inflamed ileum from patient with Crohn’s disease (**D-F**).

To define *ACE2* expression in healthy and inflamed colon, we visualized publicly available colonic sc-RNA seq data containing 366,650 cells from colonic mucosa obtained in 18 (in)active UC patients and 12 healthy individuals (Single Cell Portal, SCP 259) (**Supplementary Figure S4A-C**).^31^ As for the ileum, *ACE2* was solely expressed in colonic epithelium, mainly in a subset of enterocyte (**Figure 6A-B, Supplementary Figure S4D**). As in the ileum, the ACE2 positive colonic enterocyte cluster co-expressed *TMPRSS2* and *DPP4* (**Figure 6B**, **6E-G)**. However, in contrast to ileum, colonic *ACE2* expression was mainly restricted to enterocytes isolated form patient with active UC, while undetectable in colonic enterocytes isolated from the mucosa of healthy subjects. (**Figure 6C-D, Supplementary Figure S4E)**

**Figure 6:**
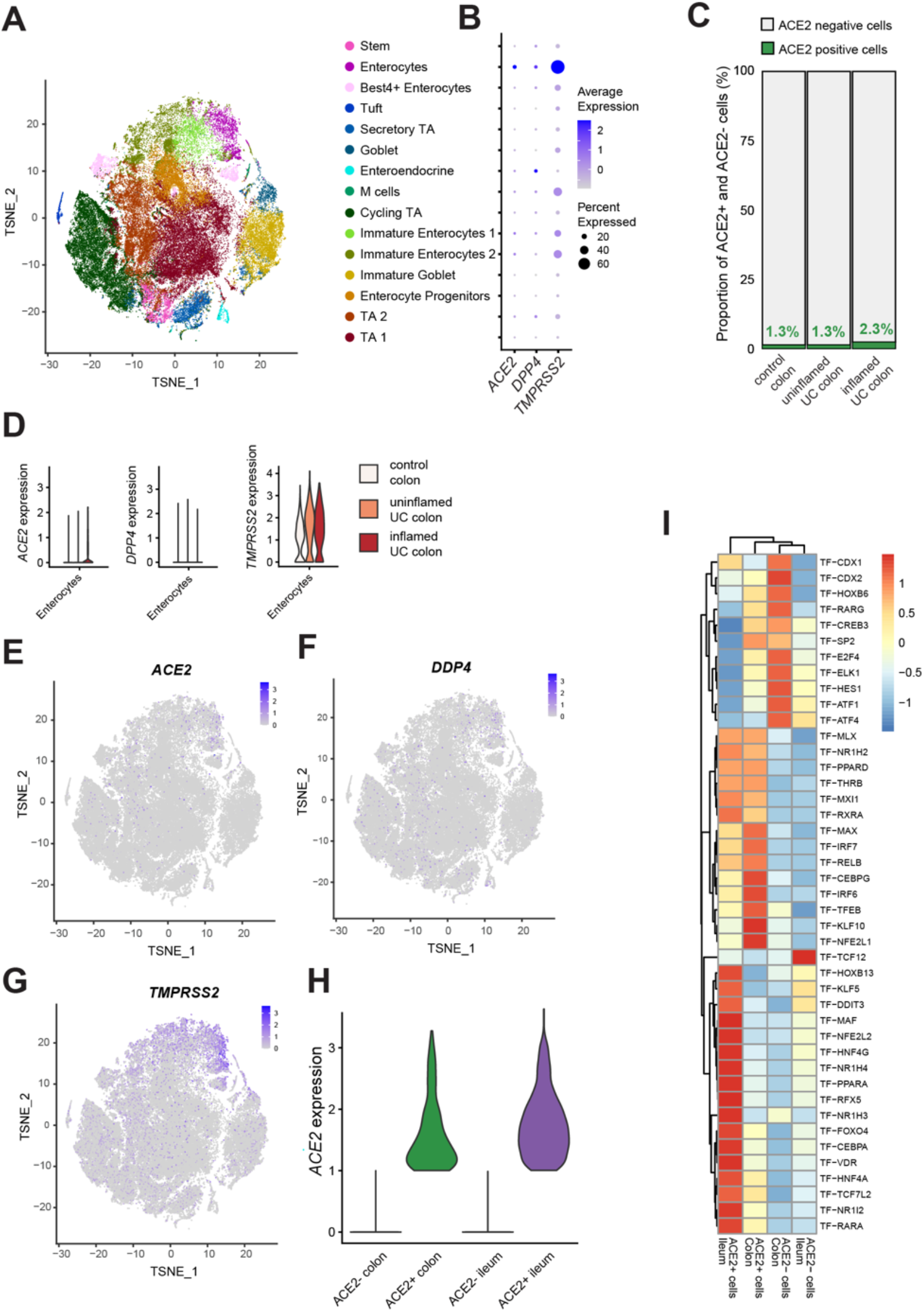
Increase colonic *ACE2* expression in the epithelial cells of patient with active UC. (**A**) t-distributed stochastic neighbor embedding (tSNE) plot showing the clustering and annotation of epithelial cells from the colon as reported in Smillie et al.^31^ (**B**) Expression of *ACE2, DPP4* and *TMPRSS2* in epithelial cell clusters of the colon. (**C**) Proportion of ACE2+ colonic epithelial cells in control, uninflamed and inflamed samples. (**D**) Expression of *ACE2, DPP4* and *TMPRSS2* in colonic enterocytes from control, uninflamed and inflamed samples. (**E-G**) Expression of *ACE2, DPP4*, and *TMPRSS2* overlaid on tSNE shown in panel A. (**H**) Expression level of *ACE2* in ACE2+ cells of colon and ileum. (**I**) Heatmap showing scaled area under the curve (AUC) values of top 15 specific and highly enriched regulons (average AUC >0.1) in ACE2+ or ACE2-compartments in integrated data of ileal and colonic single cell data identified by SCENIC analysis.

To compare expression and regulation of *ACE2* between colon and ileum, we performed an integrated analysis of epithelial cells from colon and ileum (**Supplementary Figure S5A-B).** In colonic ACE2 positive epithelial cells, *ACE2* expression was lower compared levels in ileal ACE2 positive epithelial cells (**Figure 6H**). Furthermore, using SCENIC we performed genomic regulatory networks analysis of the epithelial cells to identify specific transcription programs in *ACE2* expressing enterocytes, both in ileum and colon. As demonstrated using bulk RNA analysis, we found a relatively higher *HNF4A* regulon activation in ileal ACE2 positive cells, compared to colonic ACE2 enterocytes (**Figure 6I**). Differently, colonic ACE2 expressing enterocytes were found to have increased regulon activity of interferon responsive factors, such as IRF6 and IRF7, when compared to ileum (**Figure 6I**).

### Ileum and colon: different key regulators in ACE2 positive cells

We then asked whether particular expression patterns within ACE2 positive cells depend on the tissue and/or inflammatory state, and studied which upstream regulators were linked to these changes. When comparing expression profiles of ACE2 positive cells from inflamed CD ileum with control ileum, we found 56 differentially expressed genes (adj. p<0.05, FC>2.0). Predicted upstream regulators of these genes were HNF4A (inhibited, p=2.3E-04) and IFNγ (activated, p=5.2E-05). At the colonic level, we identified 54 differentially expressed genes in ACE2 positive cells from inflamed colon, as compared to control tissue. TNF, lipopolysaccharides, IFNγ and IL-1β were predicted as top ranked upstream regulators (activated, p≤1.9E-15)

### Inflammatory stimuli result in upregulation of *ACE2* and *TMPRSS2* in organoids from IBD patients but not from healthy individuals

Because of the clear upregulation of *ACE2* in inflamed colonic mucosa (**Figure 1A**) and the prediction of TNF as key regulator in ACE2 positive cells, we investigated the effect of an inflammatory stimulus on *ACE2* expression in an *ex vivo* organoid model. In organoids derived from controls, inflammatory stimuli did not affect *ACE2* expression (p=9.1E-01) (**Figure 7A**). Strikingly, in organoids derived from inflamed or uninflamed colonic biopsies from UC patients, addition of an inflammatory stimulus did significantly upregulate *ACE2* (FC=2.4, p=1.7E-02; FC=2.0, p=2.9E-02) (**Figure 7A**). No significant effect on *DPP4* expression could be observed (p=7.4E-02; p=7.9E-01), whereas *TMPRSS2* was significantly upregulated after inflammatory stimulation (FC=2.6, p=5.1E-14; FC=2.8, p=1.5E-30) (**Figure 7B-C**).

**Figure 7:**
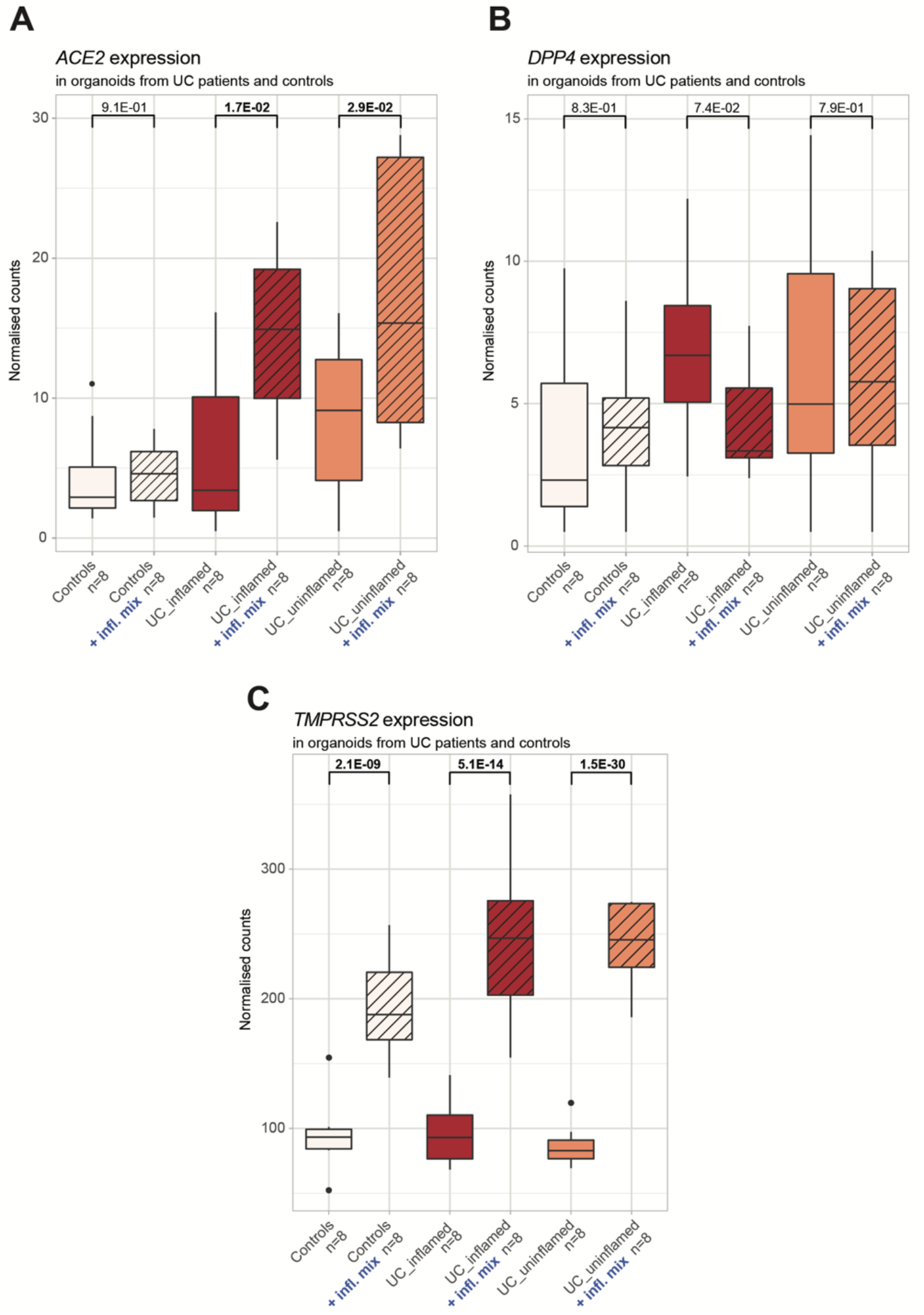
Organoid *ACE2, DPP4* and *TMPRSS2* in UC patients and controls with and without addition of an inflammatory mix. (**A**) Boxplots of organoid *ACE2* as measured by RNA sequencing (normalized counts). (**B**) Boxplots of organoid *DPP4* as measured by RNA sequencing (normalized counts). (**C**) Boxplots of organoid *TMPRSS2* as measured by RNA sequencing (normalized counts). Significant comparisons are highlighted in bold. control, non-IBD controls; IBD, inflammatory bowel disease; UC, ulcerative colitis

### Anti-TNF therapy restores colonic, but not ileal, epithelial *ACE2* dysregulation in anti-TNF responders

Given the *ex vivo* model clearly confirmed the effect of a pro-inflammatory mix, including TNF, on epithelial *ACE2* expression, we subsequently studied the effect of neutralizing TNF (through administration of infliximab) on intestinal *ACE2* expression in IBD patients with active endoscopic disease. Paired transcriptomic data, generated prior to first infliximab administration and 4-6 weeks after treatment initiation, confirmed a significant downregulation of colonic *ACE2* in endoscopic remitters, but not in non-remitters (p=1.8E-04, p=6.5E-01 respectively) (Supplementary Figure S6). In contrast, infliximab therapy did not significantly affect ileal *ACE2* expression in remitters and non-remitters (p=7.8E-02, p=2.25E-01 respectively).

## DISCUSSION

Many patients with IBD have long-term exposure to corticosteroids, thiopurines, methotrexate, small molecules and/or biological agents, classifying them as high-risk patients because of their immunosuppression. In addition, intestinal inflammation has shown to be an important risk factor for SARS-CoV-2 infection and prognosis in IBD.^14, 15, 16, 17, 18^ However, emerging evidence now suggests that IBD patients do not seem more vulnerable for COVID-19. To reconcile these observations, we investigated the role of intestinal inflammation on the potential viral intestinal entry mechanisms through bulk and single cell transcriptomics, immunofluorescence and *ex vivo* organoid cultures in patients with IBD.

In contrast to previous bulk data,^40^ we observed significant alterations in intestinal *ACE2* expression depending on the location and inflammatory state, both at tissue and single cell mRNA level, as at protein level. *ACE2* expression was limited exclusively to epithelial cells, both in ileum and colon. Hence, *ACE2* dysregulation in bulk transcriptomics as a result of massive influx of immunocytes at the site of inflammation could be excluded.

It is conceived that SARS-CoV-2 infects epithelial cells, causing cytokine and chemokine release, resulting in acute intestinal inflammation characterized by infiltration of neutrophils, macrophages and T cells,^41^ with associated shedding of faecal calprotectin and increased systemic IL-6 response,^42^ and IFN signaling.^7^ Similar to recent data,^43, 44, 45^ we found a significant downregulation of *ACE2* in inflamed ileum and a significant *ACE2* upregulation in inflamed colon. This opposing effect of inflammation on intestinal *ACE2* expression in small and large intestine was striking, which could be attributed – based on sc-RNA data – to different key transcription factors active between ileal and colonic ACE2 positive cells.

Being an IBD susceptibility locus,^46^ epithelial HNF4A plays a protective role in IBD by consolidating the epithelial barrier,^47^ especially in small intestine.^48^ As it appears to be a transcriptional sensor of inflammation,^49^ and because of its key role as transcription factor in the regulation of angiotensinogen metabolism,^50^ the decrease in *ACE2* in inflamed ileum does therefore not come as a surprise. In individuals carrying the minor *AA* genotype at the IBD HNF4A susceptibility locus, ileal *ACE2* expression was even further downregulated, without any effect on colonic *ACE2*. Of note, our sc-RNAseq data showing *ACE2* downregulation in enterocytes from inflamed CD ileum further suggest an intrinsic regulation of *ACE2*. In addition, as we observed significant correlations between enterocyte markers and ileal *ACE2* as well as an overall decrease in number of cells expressing *ACE2* in inflamed CD ileum, a loss of enterocytes might also explain lower *ACE2* levels.

Remarkably, a very recent GWAS study identified 3p21.31 as a genetic locus being associated with COVID-19-induced respiratory failure.^51^ This locus covers a cluster of 6 genes *(SLC6A20, LZTFL1, CCR9, FYCO1, CXCR6*, and *XCR1*), with the identified risk allele (i.e. worse COVID-19 outcome) being associated with increased *SCL6A20* expression. Strikingly, *SCL6A20* is known to be regulated by HNF4A.^52^

Although the colonic *ACE2* co-expression cluster in bulk tissue was also enriched for *HNF4A* as upstream transcriptional regulator, single cell data revealed that colonic *ACE2* expression seems primarily driven by interferon regulator factors. Upstream regulating analysis further supported that pro-inflammatory cytokines, including TNF, IFNγ and IL-1β contribute to colonic *ACE2* upregulation. Elevated colonic *ACE2* levels in patients with active inflammation may thus promote viral entry and, in theory, could promote COVID-19 disease severity. One can question this hypothesis as downregulated *ACE2* in inflamed ileum remains much higher than in normal and IBD colon. However, *ACE2* expression is the most abundant in the small intestine, followed by the large intestine, whereas its expression is limited in the respiratory system.^53, 54^ While there is yet no direct evidence that altered expression of intestinal *ACE2* directly impacts SARS-CoV-2 intestinal entry and tropisms to different intestinal sites,^55^ using *ex vivo* organoid models we confirmed that pro-inflammatory cytokines can upregulate colonic epithelial *ACE2* expression in IBD patients, but not in healthy individuals. Different genetic susceptibility and/or microbial composition may be responsible for difference in response to inflammatory stimuli observed in controls and IBD. Indeed, it has already been demonstrated that organoids from UC patients maintain some inherent differences as compared to non-IBD tissue,^19, 56^ presumably reflecting inherent genetic factors which could result in a more sensitive epithelium.

Being the key example of a complex immune-mediated entity where environmental and microbial factors modulate the immune response in a genetically susceptible host,^57^ the differences in *ACE2* expression upon inflammatory stimuli between colon and ileum in patients with IBD may also be attributed to differences in the intestinal microbiome. Lipopolysaccharides, comprising the wall of gram-negative bacteria, was indeed identified as one of the key drivers of the *ACE2* gene cluster in colon, but not in ileum. However, blind use of antibiotics or probiotics for COVID-19 is not recommended until a better understanding of the effect of SARS-CoV-2 on gut microbiota is obtained.^58^

In line with our findings, national and international registries suggest active IBD as a risk factor for (complicated) COVID-19.^14, 15, 16, 17, 18^ Adequate disease management, by appropriate dampening of intestinal inflammation, therefore seems key in preventing IBD patients from COVID-19. Because of the significant *ACE2* upregulation in colon, one might consider that active UC patients and/or CD patients with colonic involvement could be at higher risk for (complicated) COVID-19, compared to ileal CD. Although international registries did not yet report any outcome data split by disease location, our data would suggest an increased risk for complicated COVID-19 depending on disease location and disease activity.

Of note, several key cytokines implicated in IBD pathogenesis,^57, 59^ and also key drivers of *ACE2* colonic expression in this study, are currently under investigation as potential therapeutic targets for COVID-19, including TNF, IFNγ, IL-1β and IL-6.^60^ Although further evidence is warranted if these anti-cytokine therapies can dampen the observed cytokine storm in COVID-19, we demonstrated that anti-TNF therapy does restore intestinal *ACE2* dysregulation in a subset of IBD patients.

Although we acknowledge the lack of data on SARS-CoV-2 infected patients, a sequencing depth not enabling to look for *HNF4A* alternative splicing and isoforms with pro- and anti-inflammatory effects,^61^ and the lack of additional functional validation experiments, the replication of our findings on several levels (tissue and single cell gene expression, protein expression and *ex vivo* models) highlights the impact of our observations for the management of IBD patients in the current COVID-19 crisis. Current guidelines do not promote stopping of immunosuppressive and biological drugs in IBD patients without symptoms suggestive of COVID-19. On the contrary, immunosuppressive and biological drugs may protect against the development of severe forms of COVID-19 infection.^62^

In conclusion, using bulk and single cell transcriptomic datasets, as well as *ex vivo* organoid cultures, we demonstrated that intestinal inflammation could alter SARS-CoV-2 entry mechanisms in the intestinal epithelium, with opposing effects seen in ileum and colon. *HNF4A*, an IBD susceptibility gene and transcriptional regulator of one of the key Covid-19 GWAS loci, is an important upstream regulator of *ACE2* expression in ileal tissue. In contrast, colonic *ACE2* expression depends on interferon regulating factors and pro-inflammatory cytokines. Overall, our translational data provide further evidence for the clinical recommendation to pursue adequate disease control in patients with IBD to reduce the risk of (complicated) COVID-19.

## ABBREVIATIONS

ACE: angiotensin converting enzyme
adj: adjusted
COVID-19: coronavirus disease 2019
CD: Crohn’s disease
CRC: colorectal cancer
DPP4: dipeptidyl peptidase 4
FC: fold change
FDR: false discovery rate
HNF4A: hepatocyte nuclear factor 4 Alpha
GEM: gel beads in emulsion
IBD: inflammatory bowel disease
IFN: interferon
IL: interleukin
IPA: ingenuity pathway analysis
IQR: interquartile range
MAF: minor allele frequency
MERS: middle east respiratory syndrome
RAS: renin-angiotensinogen system
RNA: ribonucleic acid
SARS: severe acute respiratory syndrome
SCENIC: Single Cell Network Inference
TMPRSS: transmembrane protease serine
TNF: tumour necrosis factor
UC: ulcerative colitis
WGCNA: weighted gene co-expression network analysis

## SUPPLEMENTARY FIGURES

**Supplementary Figure 1: Correlation between mucosal *ACE2* levels and mucosal *HNF4A* levels in patients with IBD and controls.** Correlation between normalized *ACE2* counts and normalized *HNF4A* counts in ileal (**A**) and colonic (**B**) tissue from IBD patients and controls

**Supplementary Figure 2**: **Correlation between ileal *ACE2* levels and ileal epithelial marker gene levels in patients with IBD and controls.** Correlation between normalized ileal *ACE2* counts and normalized epithelial marker gene counts: *APOA1* (**A**), *SI* (**B**), *FABP6* (**C**) and *ENPEP* (**D**)

**Supplementary Figure 3: Ileal epithelial cell subtypes annotation and sub clusters expression of *ACE2, DPP4* and *TMPRSS2.*** (**A**) Heatmap showing the SingleR score for annotation of the 61 cell clusters of the ileum dataset. (**B**) Barplot showing the distribution of control, uninflamed and inflamed sample types in the 78,722 cells sequenced from ileum. (**C**) Expression of various epithelial sub type markers in the epithelial sub clusters of the ileum. (**D**) Expression of *ACE2, DPP4* and *TMPRSS2* in the epithelial sub clusters. E) Expression of *ACE2, DPP4* and *TMPRSS2* in the epithelial sub clusters split between control, uninflamed and inflamed sample type.

**Supplementary Figure 4: Colonic epithelial cell subtypes annotation and sub clusters expression of *ACE2, DPP4* and *TMPRSS2.*** (**A-C**) Expression of *ACE2* in the colonic epithelial cells, immune cells and stromal cells. D) Expression of epithelial subtype markers in colonic epithelial clusters. (**D**) Expression of *ACE2, DPP4* and *TMPRSS2* in colonic epithelial clusters split between sample types of control, uninflamed and inflamed sample type.

**Supplementary Figure 5: Integrated analysis of epithelial cells from colon and ileum.** (**A**) Proportion of ACE2+ cells in epithelial clusters of ileum and colon from the integrated analysis of single cell data of colon and ileum. (**B**) Contribution of the subclusters of colonic and ileal epithelial cells to ACE2+ and ACE2-compartments in colon and ileum.

**Supplementary Figure 6**: **Colonic *ACE2* expression in IBD patients prior and after anti-TNF therapy.** Boxplots of normalized log2 transformed *ACE2* expression levels prior to and 4-6 weeks after infliximab therapy in colonic mucosa of IBD patients, split by endoscopic remission.

Endoscopic remission in UC: Mayo endoscopic sub-score 0-1; endoscopic remission in CD: complete absence of ulcerations

## TABLES

**Supplementary Table 1: Demographics of all included patients**

**Supplementary Table 2: Genes within ileal *ACE2*-coexpression module**

**Supplementary Table 3: Genes within colonic *ACE2*-coexpression module**

## ACKNOWLEDGEMENTS

The authors would like to thank Vera Ballet, Helene Blevi, Sophie Organe, Nooshin Ardeshir Davani and Tamara Coopmans for an excellent job in maintaining the Biobank database; André D’Hoore and Gabriele Bislenghi (Department of Abdominal Surgery, University Hospitals Leuven, Belgium) for the resection specimens; Gabriele Dragoni and Brecht Creyns (Translational Research in GI disorders KU Leuven, Belgium) and Gert De Hertogh (Laboratory of Morphology and Molecular Pathology, University Hospitals Leuven, KU Leuven, Leuven, Belgium) for the processing of the resection specimens; Birgit Weynand, Lukas Marcelis and Matthias Van Haele (Laboratory of Morphology and Molecular Pathology, University Hospitals Leuven, KU Leuven, Leuven, Belgium) for the ACE2 antibody; David Carbonez, Kristine Stepnayan, Nina Dedoncker, Vanessa Brys, Jens Van Bouwel, Wim Meert, Alvaro Cortes Calabuig and Wouter Bossuyt (Genomics Core Facility, University Hospitals Leuven, Belgium) for the technical assistance with the RNA-sequencing library preparation and sequencing. Immunostainings were recorded on a Zeiss LSM 780 – SP Mai Tai HP DS (Cell and Tissue Imaging Cluster (CIC), supported by Hercules AKUL/11/37 and FWO G.0929.15 to Pieter Vanden Berghe, KU Leuven, Leuven, Belgium.

## AUTHOR CONTRIBUTIONS

BV and SV contributed equally and share the first authorship. GM and SV contributed equally and shared the senior authorship. BV: study design, data acquisition and interpretation, statistical analysis and drafting of the manuscript. SaV: study design, data acquisition and interpretation, statistical analysis and drafting of the manuscript. SAR: data acquisition and interpretation (single cell RNA), statistical analysis and critical revision of the manuscript. BJK: data acquisition and interpretation (single cell RNA and immunostainings) and critical revision of the manuscript. KA: data acquisition and interpretation (organoid data), statistical analysis and critical revision of the manuscript. IC: data acquisition (genetics) and critical revision of the manuscript. JS: data interpretation and critical revision of the manuscript. MF: data interpretation and critical revision of the manuscript. GM: supervision, data acquisition and interpretation, critical revision of the manuscript. SV: study design, supervision, data interpretation and critical revision of the manuscript. All authors agreed on the final manuscript.

Guarantor of the manuscript: Séverine Vermeire.

## CONFLICTS OF INTEREST

B Verstockt reports financial support for research from Pfizer; lecture fees from Abbvie, Ferring, Takeda Pharmaceuticals, Janssen and R Biopharm; consultancy fees from Janssen and Sandoz.

J Sabino reports lecture fees from Abbvie, Takeda, Janssen and Nestle Health Sciences.

M Ferrante reports financial support for research: Amgen, Biogen, Janssen, Pfizer, Takeda, Consultancy: Abbvie, Boehringer-Ingelheim, MSD, Pfizer, Sandoz, Takeda and Thermo Fisher; Speakers fee: Abbvie, Amgen, Biogen, Boehringer-Ingelheim, Falk, Ferring, Janssen, Lamepro, MSD, Mylan, Pfizer, Sandoz, and Takeda.

G Matteoli received financial support for research from DSM Nutritional Products, Karyopharm Therapeutics and Janssen.

S Vermeire reports financial support for research: MSD, AbbVie, Takeda, Pfizer, J&J; Lecture fee(s): MSD, AbbVie, Takeda, Ferring, Centocor, Hospira, Pfizer, J&J, Genentech/Roche; Consultancy: MSD, AbbVie, Takeda, Ferring, Centocor, Hospira, Pfizer, J&J, Genentech/Roche, Celgene, Mundipharma, Celltrion, SecondGenome, Prometheus, Shire, Prodigest, Gilead, Galapagos.

S Verstockt, S Abdu Rahiman, BJ Ke, K Arnouts and I Cleynen declare no conflicts of interest.

## FUNDING

K Arnauts is a doctoral fellow and S Vermeire and M Ferrante are Senior Clinical Investigators of the Research Foundation Flanders (FWO), Belgium. G Matteoli laboratory is supported by a FWO grant (G.0D83.17N), a grant from the International Organization for the Study of Inflammatory Bowel Diseases (IOIBD), a grant from the European Crohn’s and Colitis Organization (ECCO) and grants from the KU Leuven Internal Funds (C12/15/016 and C14/17/097). S Vermeire and G Matteoli are funded by Strategic Basic Research FWO grant (S008419N).

