## Supplementaries for "Intestinal receptor of SARS-CoV-2 in inflamed IBD tissue is downregulated by HNF4A in ileum and upregulated by interferon regulating factors in colon"

#### Supplementary Figure S1

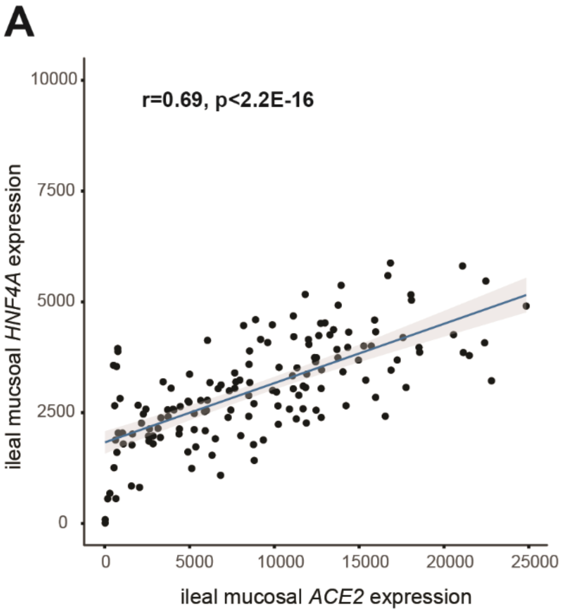

**Supplementary Figure S2**

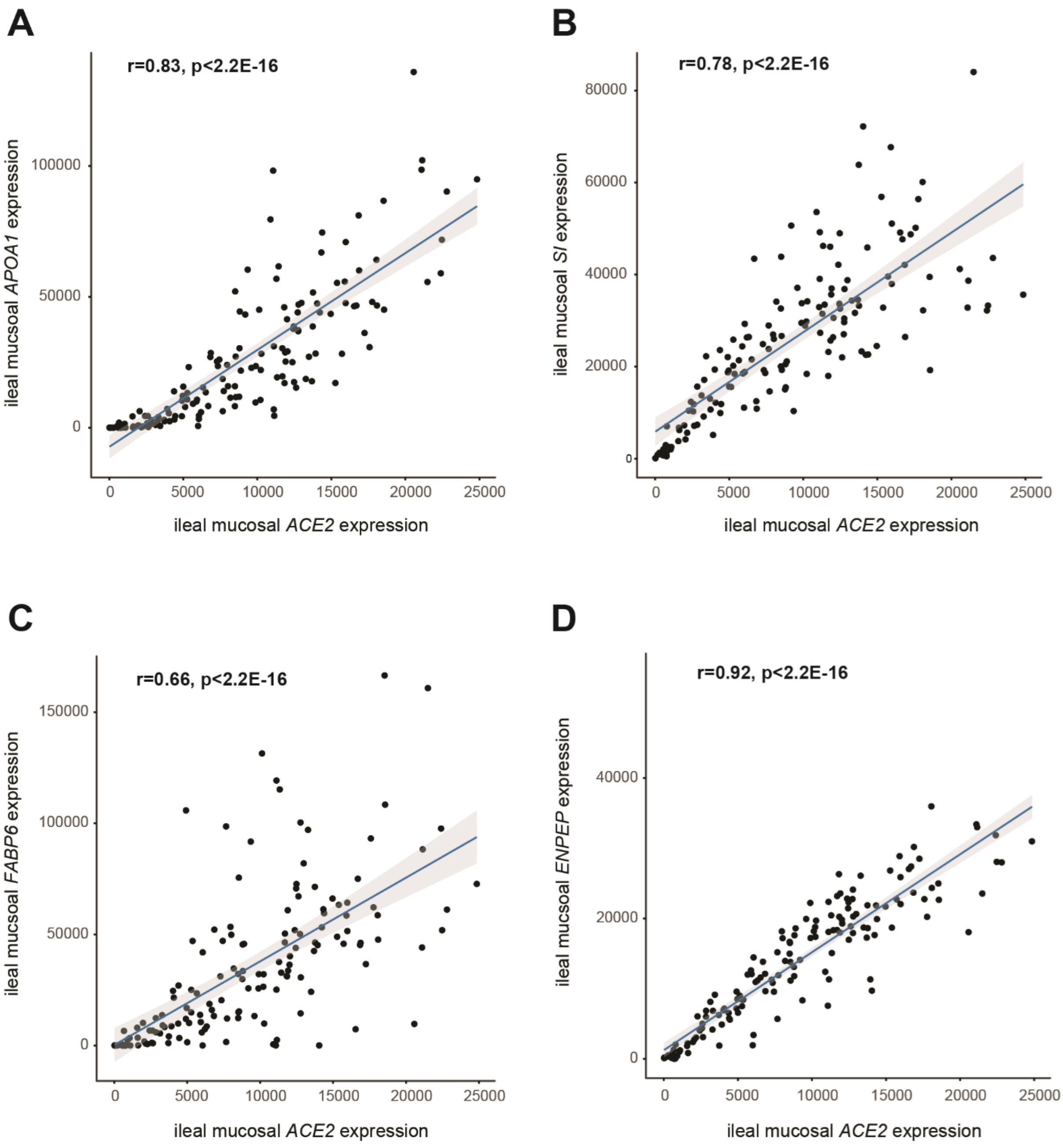

#### Supplementary Figure S3

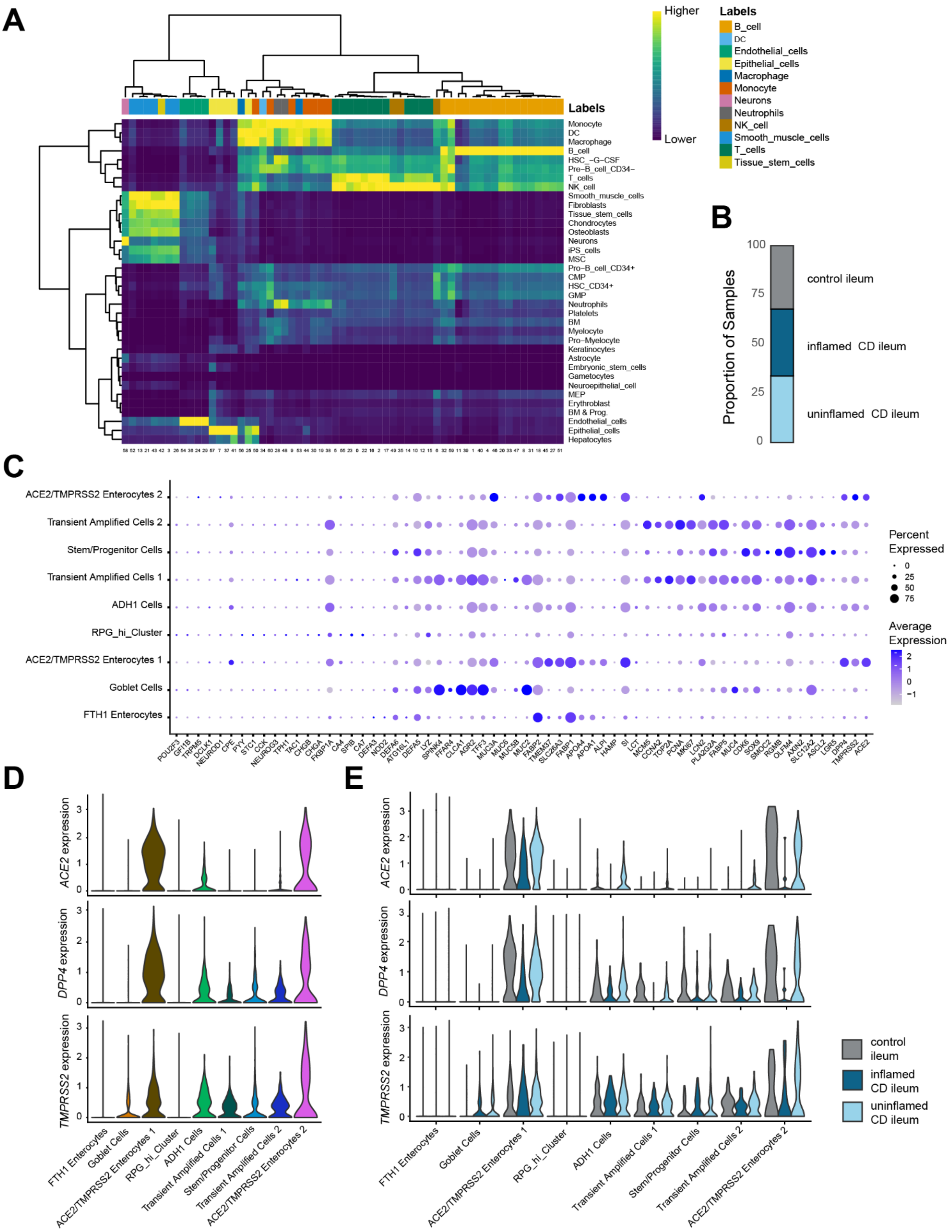

### Supplementary Figure S4

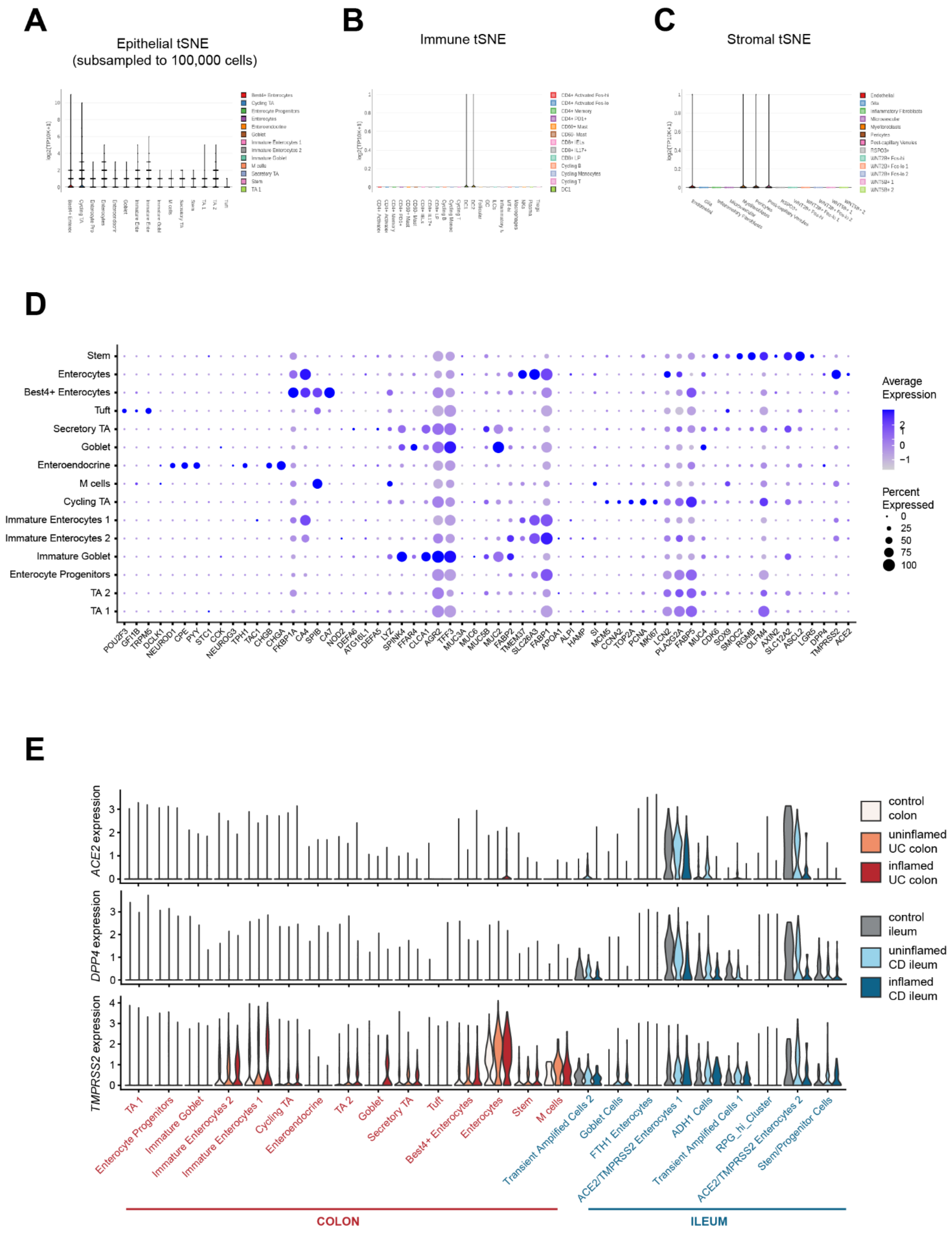

#### Supplementary Figure S5

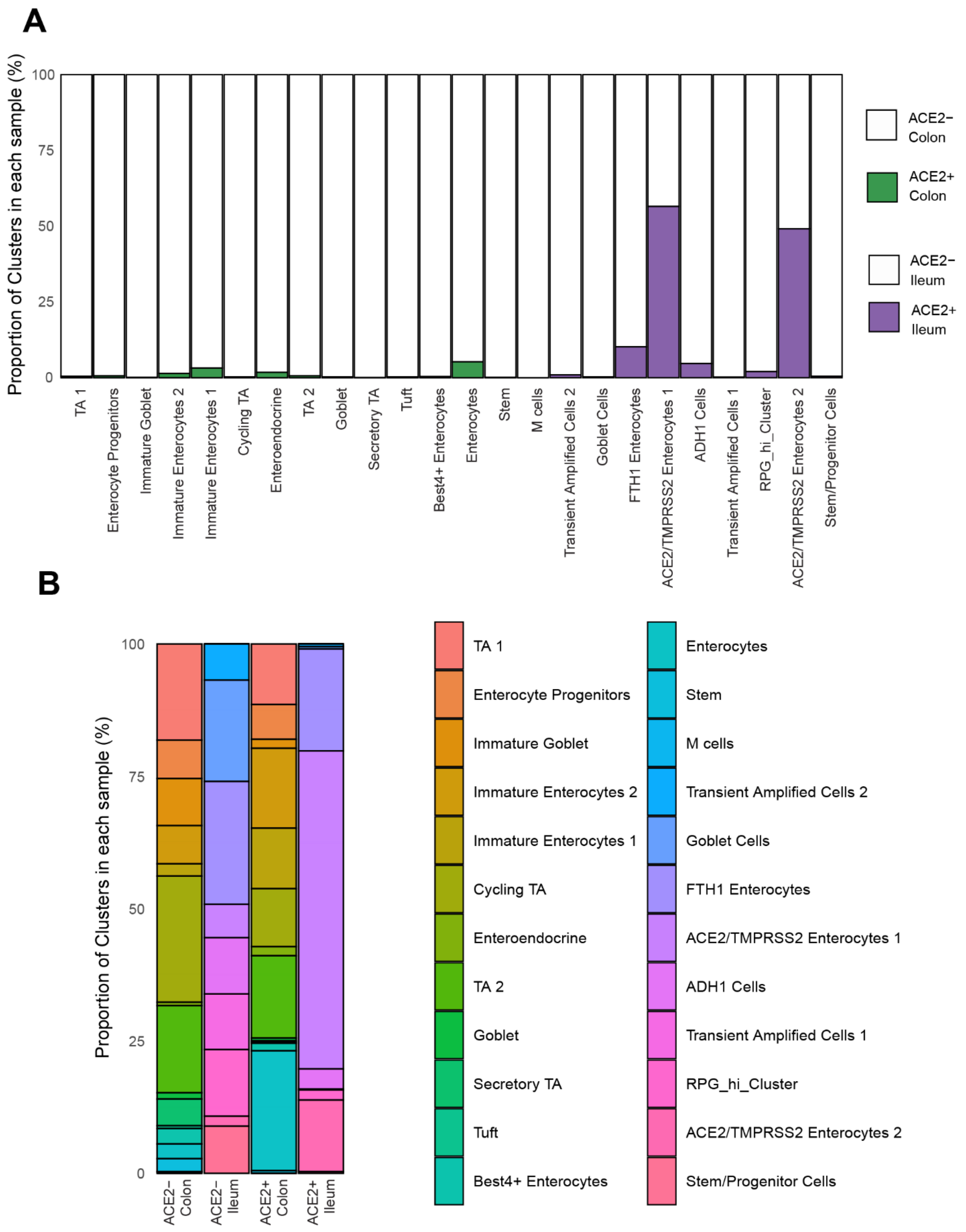

#### Supplementary Figure S6

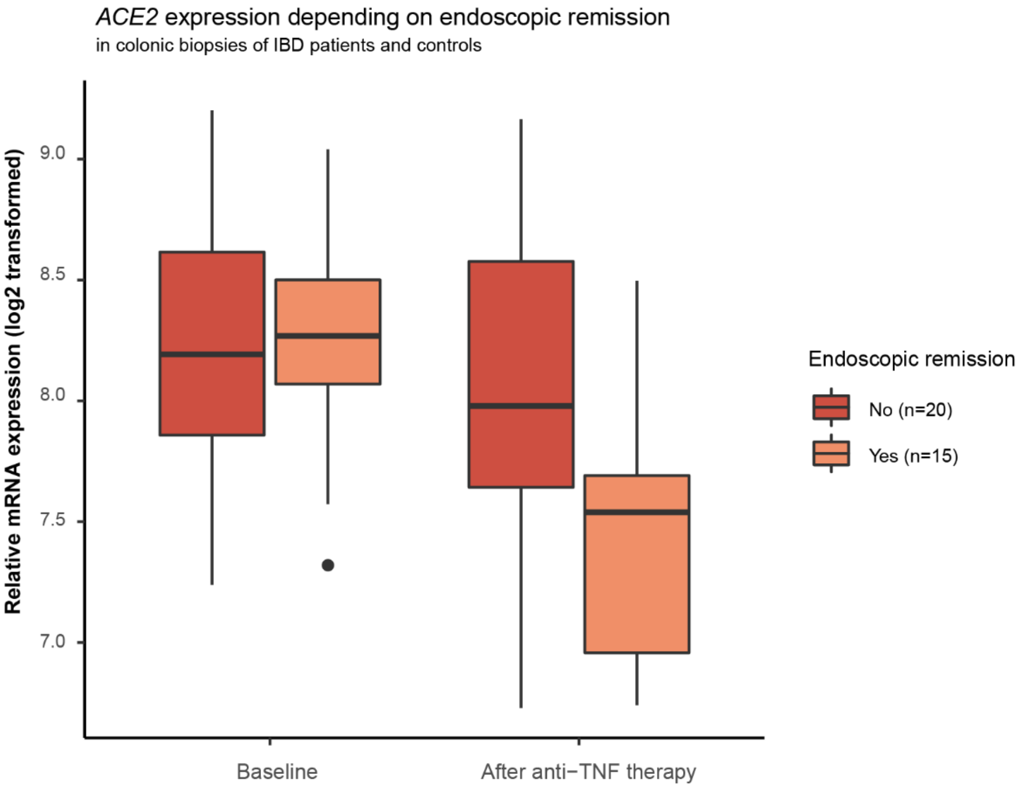

**Supplementary Table 1**: Baseline demographics from included patients

| | **Bulk transcriptomics** | | **Single cell transcriptomics** | | **Organoids** | |
|---|---|---|---|---|---|---|
| | IBD (n = 351) | non-IBD controls (n = 51) | IBD (n = 6 ) | non-IBD controls (n= 5) | IBD (n = 8) | non-IBD controls (n =8) |
| **Disease** | | | | | | |
| - **Crohn's disease** | 193 (55.0) | N.A. | 6 (100.0) | N.A. | 0 (0.0) | N.A. |
| - **Ulcerative colitis** | 158 (45.0) | | 0 (0.0) | | 8 (100.0) | |
| **Women, *n* (%)** | 183 (52.1) | 34 (66.7) | 5 (83.3) | 1 (20.0) | 4 (50.0) | 5 (62.5) |
| **Age, *years,* median [IQR]** | 41.8 (26.7 – 53.0) | 57.0 (41.0 – 63.0) | 42.7 (38.5 – 49.5) | 71.0 (52.5 – 73.0) | 41.3 (35.9 – 46.5) | 45.0 (30.5 – 55.9) |
| **Disease duration, *years*, median [IQR]** | 7.9 (2.0 – 16.8) | N.A. | 13.4 (5.8 – 22.1) | N.A | 10.5 (7.8 – 12.7) | N.A. |
| **Disease location, n (%)** | | | | | | |
| - Ileal (L1) | 41 (21.3) | N.A. | 3 (50.0) | N.A. | N.A. | N.A. |
| - Colonic (L2) | 23 (11.9) | | 0 (0.0) | | | |
| - Ileocolonic (L3) | 129 (66.8) | | 3 (50.0) | | | |
| - Upper GI modifier (+L4) | 57 (29.5) | | 0 (0.0) | | | |
| - Proctitis (E1) | 13 (8.2) | N.A. | N.A. | N.A. | 1 (12.5) | N.A. |
| - Left-sided colitis (E2) | 85 (53.8) | | | | 1 (12.5) | |
| - Extensive colitis (E3) | 60 (38.0) | | | | 6 (75.0) | |
| **Disease behaviour, n (%)** | | | | | | |
| - Inflammatory (B1) | 95 (49.2) | N.A. | 0 (0.0) | N.A. | N.A. | N.A. |
| - Fibrostenotic (B2) | 55 (28.5) | | 6 (100.0) | | | |
| - Penetrating (B3) | 43 (22.3) | | 0 (0.0) | | | |
| - Perianal modifier (p) | 32 (16.6) | | 1 (16.7) | | | |

n = number of patients; IQR = interquartile range; N.A. = not applicable

#### Supplementary Table 2: Genes within ileal *ACE2*-coexpression module

| | Genes |
|---|---|
| **Ileal *ACE2* co-expression module "blue"** | A1CF, AADAC, AARS2, ABAT, ABCA5, ABCA8, ABCB1, ABCB7, ABCC2, ABCC8, ABCG2, ABHD15, ABHD6, AC009403.2, AC022431.2, AC079602.1, ACACB, ACAD11, ACADM, ACBD5, ACE, ACE2, ACLY, ACOT12, ACOT4, ACOX1, ACOX2, ACRBP, ACSF2, ACSS1, ACVR2A, ADAMDEC1, ADAMTS17, ADCY2, ADD1, ADD3, ADIPOR2, ADORA3, ADPRM, ADRA1B, ADRB1, AFG3L2, AGBL2, AGFG1, AGGF1, AGMO, AGPAT3, AGPAT9, AGXT2, AHCYL1, AK3, AKAP6, AKTIP, AL009178.1, ALAD, ALDH18A1, ALDH6A1, ALDH9A1, ALDOC, ALKBH1, ALS2, AMFR, AMOT, ANAPC16, ANKRA2, ANKRD27, ANKRD46, ANO10, ANO5, ANO6, AOC1, AP000783.1, AP1S3, APOA1, APOA4, APOB, APOM, AQP1, AQP3, AQP7, AREL1, ARHGAP18, ARHGAP32, ARHGEF11, ARHGEF26, ARHGEF38, ARHGEF7, ARL5A, ARMC1, ARMCX5, ARRDC4, ASAH2, ASAH2C, ASB8, ASH2L, ASPA, ASTE1, ASTN2, ASXL3, ATG12, ATG16L1, ATG5, ATL1, ATP1A1, ATP6V0A2, ATRN, ATXN7L1, AUH, AUTS2, AZIN1, B3GALT2, B3GNT5, B4GALT4, B4GALT6, BACH1, BAG4, BAHD1, BAIAP3, BBS12, BCAR3, BCL2L10, BCL2L15, BCO2, BCORL1, BEND7, BHLHB9, BIRC2, BLNK, BLOC1S5, BMF, BMP3, BMP5, BMX, BRAP, BRD1, BRD7, BRI3BP, BRINP3, BST1, BTNL2, BUD13, C10orf32, C10orf76, C14orf28, C16orf72, C16orf87, C17orf103, C17orf78, C17orf80, C1D, C1orf106, C1orf115, C1orf131, C1orf168, C20orf202, C21orf49, C21orf88, C22orf29, C2orf68, C2orf69, C2orf88, C4orf29, C5orf15, C6, C6orf123, C9orf24, C9orf40, C9orf78, CA10, CABP7, CACNA1D, CACNA2D4, CACNB2, CAMKK2, CAPN13, CAPZA2, CASC5, CASK, CASP6, CAT, CATSPERG, CBS, CBWD6, CCDC126, CCDC127, CCDC152, CCDC174, CCDC28A, CCDC68, CCDC91, CCL14, CCNDBP1, CCNG1, CCNY, CCR9, CCSER1, CD160, CD163L1, CD244, CD36, CD58, CD8B, CDC14A, CDC14B, CDC42EP4, CDH16, CDHR1, CDHR2, CDHR3, CDK20, CDK3, CDK8, CDKN2AIP, CDKN2B, CDR2, CDS1, CDYL2, CEACAM18, CEBPZ-AS1, CELF3, CENPJ, CEP76, CERK, CERKL, CERS6, CGRRF1, CHAD, CHCHD7, CHDH, CHGB, CHMP4C, CHN2, CHPT1, CHRAC1, CHRFAM7A, CHRM4, CHRNA7, CHST13, CHST5, CHST6, CIAO1, CIR1, CITED2, CKMT2, CLCA2, CLCN1, CLCN5, CLDN8, CLEC4F, CLIC5, CLN5, CLPTM1, CLPX, CLSTN2, CMBL, CMPK1, CMTR2, CNGA1, CNGA3, CNIH1, CNKSR3, CNNM3, CNOT7, CNOT8, CNTFR, CNTNAP2, COBL, COG5, COL17A1, COL2A1, COLEC11, COPA, COPS2, COPS5, COX11, COX20, CPEB2, CPEB3, CPLX2, CPO, CPOX, CPPED1, CPQ, CPS1, CREB3L2, CRIP1, CRIPT, CRNKL1, CROT, CRY2, CRYBA2, CRYZ, CSNK1G3, CSRP2BP, CTAGE5, CTC-534A2.2, CTH, CTPS2, CTSO, CUBN, CXXC4, CYB561D1, CYBRD1, CYP2B6, CYP2C19, CYP2C8, CYP2J2, CYP2R1, CYP2U1, CYP3A4, CYP3A7, CYP4F2, CYP4F3, CYP4V2, CYTH3, DAB1, DACH1, DAPK1, DAZAP2, DCAF8, DCUN1D2, DDB1, DDHD2, DDO, DDX19B, DDX27, DDX52, DDX58, DEFA5, DEFA6, DENND5B, DEPDC7, DEPTOR, DERA, DET1, DFFB, DGKD, DHFR, DHFRL1, DHRS4-AS1, DHRS7, DIP2C, DIRAS2, DISP1, DNAAF2, DNAH1, DNAH10, DNAJA2, DNAJB13, DNAJB7, DNAJC19, DNAJC22, DNAJC25, DNASE1, DOCK5, DPEP1, DPH3, DPP10, DPP4, DRAM2, DSCR3, DUSP11, DUSP19, DUSP3, EAPP, ECM2, EDN1, EDN3, EFTUD1, EGLN1, EIF1B, EIF4A2, EIF4EBP2, EIF5, ELMOD2, ELOVL7, EMB, EMC2, ENHO, ENPEP, ENPP3, ENPP6, ENTPD7, EPB41L3, EPDR1, EPHX1, EPT1, ERAP1, ERBB2, ESCO2, ESPL1, ESRRG, ETFDH, EXOC6B, EXT1, F10, FABP6, FADS6, FAF1, FAM102A, FAM110C, FAM134B, FAM13A, FAM149B1, FAM151A, FAM160B1, FAM168A, FAM189A1, FAM193A, FAM200B, FAM213A, FAM35A, FAM3C, FAM46A, FAM47E, FAM53B, FAM83F, FAM8A1, FANCC, FASTKD3, FASTKD5, FBXL20, FBXO22, FBXO25, FBXO27, FBXO31, FBXO48, FBXO8, FCER1A, FEZ2, FGF19, FGF9, FGFR1OP, FGGY, FHDC1, FLCN, FLVCR1, FLVCR2, FMO1, FMO4, FMO5, FNTA, FOSL2, FOXO4, FOXP2, FREM1, FRG1B, FRK, FRMD3, FSD1L, FUCA2, FZD1, FZD7, G6PC, GABARAPL1, GABRA2, GABRA4, GALNT1, GALNT13, GALNT14, GARNL3, GATA6, GATAD1, GATC, GATM, GATS, GBF1, GBP3, GCG, GCLC, GCLM, GCNT1, GCNT2, GCNT4, GCOM1, GDA, GDPD1, GDPD2, GFOD1, GFRA2, GGNBP2, GIN1, GIT1, GK, GLB1L, GLOD4, GLP2R, GLS, GLUD1, GLUD2, GNG12, GNG4, GNS, GOLGA7, GPATCH2, GPN2, GPR112, GPR128, GPR158, GPRIN2, GRAMD1B, GRAMD1C, GRIA4, GRIK2, GRIN2A, GRIP1, GSTA1, GSTA2, GSTA5, GTF2H1, GUCA2A, GUCA2B, GUCY2C, GZMA, H2AFV, HACE1, HAPLN1, HBP1, HDAC5, HHIP, HHLA2, HINFP, HIRIP3, HIST1H3E, HIST4H4, HLF, HNF4G, HNRNPH2, HNRNPH3, HOOK1, HOXA2, HOXB7, HOXC11, HOXD1, HPGD, HPGDS, HPS3, HRH1, HS2ST1, HSD17B11, HSD3B1, HSDL2, HTR1D, HTR4, HYKK, ICMT, ID2, IER3IP1, IFIT1, IFIT2, IFNLR1, IGIP, IGSF23, IL13RA1, IL17RB, IL6R, IL7, IMMP2L, ING1, INSIG2, INSM1, IPMK, IRF2BP2, IRS2, ISCA1, ISL1, ISOC1, IST1, ISX, ITGAE, ITGB8, ITLN2, ITM2B, IYD, KAT2B, KB-1507C5.2, KBTBD11, KBTBD12, KBTBD3, KCNG1, KCNH6, KCNJ13, KCNJ16, KCNJ3, KCNQ3, KCTD3, KDM3A, KDM4A, KDM6A, KDM8, KDSR, KIAA0247, KIAA1161, KIAA1211, KIAA1211L, KIAA1467, KIAA1683, KIAA1919, KIAA1958, KIF13A, KIF3B, KL, KLB, KLHDC2, KLHL13, KLHL23, KLHL34, KLHL7, KLKB1, L3MBTL4, LACC1, LACE1, LACTB2, LAMA1, LAMB4, LARGE, LCLAT1, LCN15, LCORL, LEAP2, LGALS2, LGALSL, LGMN, LHFPL2, LHX4, LIN7A, LIPA, LIPE, LIPT1, LMBRD1, LMO7, LMX1A, LPAR5, LPGAT1, LPIN2, LRAT, LRP1, LRP4, LRRC16A, LRRC28, LRRC3, LRRC40, LRRC8D, LRRN3, LUZP1, LY75, LYPLAL1, LYRM5, LZIC, MACC1, MAF, MALRD1, MAN1A1, MAN2A1, MAOA, MAOB, MAP3K13, MAP7D2, MAPK6, MAPK8, MAPKBP1, MAST2, MAT2B, MATN2, MB21D2, MBIP, MBP, MCOLN3, MCUR1, ME2, MED17, MED29, MED4, MEP1A, MEP1B, MEPCE, METTL20, METTL24, MFI2, MFN2, MFSD8, MGAM, MGAT3, MGAT4A, MGME1, MIA2, MIA3, MIEF1, MINPP1, MKNK2, MLIP, MLK4, MME, MMEL1, MMP24, MOAP1, MOB3B, MOCS1, MPP5, MPP6, MPZL3, MRO, MS4A10, MS4A8, MSMO1, MT1A, MT1F, MT1G, MT1H, MT1HL1, MT1M, MT2A, MTERFD3, MTFR1, MTMR10, MTMR11, MTMR4, MTO1, MTRF1L, MTTP, MUC17, MYH3, MYO5B, MYO6, MYOM2, MYOM3, NAA40, NAALADL1, NAGA, NAPB, NAPEPLD, NAT8, NBAS, NBPF4, NCKAP5, NCOA4, NDFIP1, NDRG1, NDRG3, NDST1, NDUFA5, NECAP1, NEDD4, NEDD9, NELL2, NEURL1B, NEUROD1, NFE2L2, NFIB, NFRKB, NFS1, NHSL1, NMT1, NMT2, NMUR1, NOA1, NPY, NR1H4, NR1I3, NR5A2, NR6A1, NRDE2, NRK, NSL1, NT5DC1, NTS, NUAK2, NUDT13, NUDT16, NUDT19, NUDT7, NUDT9, NUMA1, NUMB, OAT, OGDH, OGG1, OPA3, ORC2, OSBPL1A, OSBPL6, OSR2, OXNAD1, P2RY2, P2RY4, PAAF1, PALB2, PALM3, PANK1, PAPD5, PAQR3, PAQR5, PARD6B, PARG, PARK2, PARP16, PARP6, PAX6, PAXBP1, PBLD, PBXIP1, PCBD2, PCK1, PCSK2, PCSK5, PCYOX1, PDGFC, PDK2, PDK3, PDK4, PDXDC1, PDZK1, PELI2, PEX2, PEX26, PEX3, PEX5, PFKFB4, PFN2, PGAP1, PGAP2, PGAP3, PGBD2, PGPEP1, PGRMC1, PGRMC2, PHEX, PHF7, PHKA2, PHLPP2, PHYH, PHYHIPL, PI4K2B, PICALM, PIGC, PIGH, PIK3C2G, PITX2, PIWIL2, PIWIL4, PKLR, PLA2G4C, PLA2R1, PLAGL2, PLB1, PLCH2, PLCXD2, PLCXD3, PLD1, PLEKHA3, PLEKHA7, PLEKHG6, PLOD2, PLRG1, PLS1, PMP22, PMPCB, PNPLA1, POF1B, POFUT1, POLE, POMGNT1, POMT1, |

| | |
|---|---|
| | *PON3, POU2F3, PPAPDC2, PPFIBP2, PPID, PPL, PPM1A, PPM1D, PPP1CC, PPP1R2, PPP1R3D, PPP2R2C, PRDX3, PRKAB2, PRKCA, PRKG2, PRPF39, PRR26, PRTFDC1, PRUNE, PSD4, PSEN1, PTGDR2, PTGR1, PTGR2, PTK2, PTK2B, PTPRR, PTTG1IP, PUS10, PVRL3, PXMP4, PYGO2, PYURF, PYY, QPRT, R3HCC1L, R3HDM2, RAB11FIP4, RAB22A, RAB6A, RAB7L1, RABGAP1L, RAF1, RAG1, RALGPS1, RARS2, RASL10B, RBBP9, RCOR3, RDH10, RDH5, REEP3, REPS1, RERE, REV1, RFX6, RGS13, RHOBTB2, RHOT1, RHOU, RIMBP2, RIMS2, RIN2, RIOK3, RIPK1, RMDN1, RMND1, RNF11, RNF125, RNF128, RNF14, RNF141, RNF157, RNF170, RNF185, RNF19A, RNFT1, RORC, RP11-451M19.3, RP11-644F5.10, RPA1, RPA3-AS1, RPP14, RPRD1B, RRAS2, RRNAD1, RSPRY1, RTN4, RUNDC3B, RXRA, SAP30L, SARM1, SC5D, SCAPER, SCG2, SCGN, SCIN, SCMH1, SCML1, SCN4B, SCN9A, SCRN3, SCUBE1, SDHD, SDR42E1, SEC16B, SEC22C, SEC23A, SEC24A, SEC24D, SEC31A, SEMA3E, SEMA3G, SEMA6D, SENP2, SEPP1, SEPSECS, SERPINA7, SESN1, SETD3, SETD9, SEZ6L, SFT2D2, SGK1, SGK223, SGK3, SGMS1, SGPL1, SH2D1B, SH2D6, SH2D7, SH3YL1, SHBG, SHISA2, SHMT1, SI, SIAH1, SIGLEC15, SIMC1, SIPA1L3, SIRT4, SKP1, SLC10A2, SLC13A1, SLC14A2, SLC15A1, SLC16A10, SLC16A4, SLC17A4, SLC17A7, SLC17A8, SLC19A3, SLC1A1, SLC1A7, SLC22A4, SLC22A5, SLC23A1, SLC23A3, SLC25A15, SLC25A16, SLC25A42, SLC25A51, SLC27A2, SLC28A1, SLC28A2, SLC2A2, SLC2A9, SLC30A1, SLC30A10, SLC30A4, SLC30A8, SLC34A3, SLC35E2B, SLC35F2, SLC35G1, SLC36A1, SLC38A11, SLC39A14, SLC39A9, SLC3A1, SLC41A2, SLC46A1, SLC46A3, SLC4A7, SLC5A11, SLC5A12, SLC5A4, SLC5A9, SLC6A19, SLC6A20, SLC6A4, SLC7A7, SLC7A9, SLC8A1, SLC8B1, SLC9A6, SLCO1C1, SLCO2B1, SLU7, SMAD3, SMAD7, SMARCD1, SMG5, SMIM19, SMLR1, SMURF2, SMYD4, SNAP23, SNRK, SNW1, SNX1, SNX14, SNX2, SNX24, SNX33, SNX4, SNX5, SOAT2, SOCS6, SORBS3, SOSTDC1, SOWAHA, SOX6, SP1, SPAG5, SPAG9, SPATA25, SPATA5L1, SPATA6L, SPPL3, SPTLC3, SRD5A1, SSTR1, SSX2IP, ST13, ST18, STAMBP, STAT6, STAU2, STK17B, STK35, STRADB, STRIP1, STX7, SUCLA2, SUCO, SUGCT, SULT1B1, SULT2A1, SUSD2, SVIP, SYNPO, TADA1, TADA2B, TAF4, TAF7, TAPT1, TAT, TBC1D13, TBC1D15, TBC1D23, TBC1D32, TBC1D8B, TBX3, TCN2, TCP11L1, TDP2, TEC, TEF, TEP1, TESK2, TEX261, TFDP2, THNSL2, THRA, TIAM2, TIGD2, TIGD3, TINAG, TIPARP, TIRAP, TJP2, TLR3, TM4SF4, TM9SF2, TMBIM6, TMCC3, TMCO1, TMEM116, TMEM117, TMEM128, TMEM133, TMEM135, TMEM140, TMEM144, TMEM14A, TMEM164, TMEM181, TMEM19, TMEM220, TMEM229A, TMEM236, TMEM252, TMEM41A, TMEM41B, TMEM52, TMEM57, TMEM59, TMEM64, TMEM80, TMEM98, TMEM9B, TMIGD1, TMTC4, TNFRSF10B, TNFRSF11A, TNFSF10, TNFSF15, TNIK, TNNC2, TOB1, TOB2, TOM1L1, TOR1AIP2, TOX4, TPH1, TPP1, TPPP, TRAF6, TRAK1, TRANK1, TRDMT1, TREH, TRIM10, TRIM32, TRIM35, TRIM36, TRIM4, TRMT10A, TRMT11, TRMT61B, TRPM5, TRPV3, TSHZ1, TSNARE1, TSPAN12, TSPAN3, TSSK6, TTC17, TTC19, TTC31, TTLL6, TTR, TUFT1, TULP4, TYW5, UACA, UBA5, UBE2B, UBE2D4, UBE2G1, UBE4B, UBQLN1, UBR2, UBTD2, UCHL5, UCK1, UCK2, UCN3, UGCG, UGT1A10, UGT1A4, UGT1A6, UGT2A3, UGT2B15, UGT2B17, UGT3A1, UMODL1, UNC5D, UNC93A, USH1G, USP3, USP42, VAMP7, VAT1, VAV2, VDR, VIL1, VIPR1, VPS35, VPS36, VPS37A, VSIG10, VSIG10L, VTA1, VTI1B, VWA5A, VWA5B2, VWA7, WBP4, WDR20, WDR49, WDR59, WDR60, WDTC1, WEE1, WFIKKN1, WIPF3, WRB, WWC1, XKRX, XPC, XPNPEP2, YIPF5, ZBED1, ZC3H12B, ZC3H15, ZC4H2, ZCCHC14, ZCCHC4, ZCCHC9, ZDHHC11, ZDHHC11B, ZDHHC6, ZDHHC9, ZFAND1, ZFAND6, ZFP41, ZFYVE1, ZG16, ZHX3, ZNF124, ZNF16, ZNF180, ZNF181, ZNF19, ZNF223, ZNF224, ZNF233, ZNF250, ZNF274, ZNF3, ZNF350, ZNF384, ZNF395, ZNF488, ZNF493, ZNF512B, ZNF528, ZNF554, ZNF577, ZNF600, ZNF606, ZNF664, ZNF689, ZNF763, ZNF782, ZNF799, ZNF821, ZNF827, ZNF836, ZP2, ZSCAN16, ZSCAN20, ZSCAN30, ZSWIM3, ZZEF1* |

**Supplementary Table 3: Genes within colonic *ACE2*-coexpression module**

| | **Genes** |
|---|---|
| **Colonic *ACE2* co-expression module "green"** | *ABCD4, ABCG1, ABHD12, ABHD4, ABLIM1, ABLIM2, ABTB2, AC009403.2, ACAA1, ACADS, ACAT2, ACE2, ACOT1, ACOT11, ACOT2, ACOT4, ACOT8, ACTN4, ACTR10, ACVRL1, ADIRF, ADORA2B, ADRA2A, AGPAT2, AGPAT3, AGPAT9, AGR3, AIM1L, AK1, AK7, AKR1B10, AKR1B15, AKR1C2, AKR1C3, ALDH1L1, ALDH2, ALDH3A2, ALPPL2, AMFR, AMN, ANKRD54, ANKS4B, AOC1, AP003068.23, AP1M2, AP2A2, AP4M1, APLP2, APOBEC1, APOBR, APP, ARAF, ARHGAP39, ARHGEF16, ARHGEF18, ARHGEF35, ARHGEF5, ARL4A, ARL6IP5, ARRDC1, ARTN, ATG16L1, ATG4D, ATG9A, ATP10B, ATP1B1, ATP1B3, ATP6V0A1, ATP6V0D1, ATP6V1D, ATP7B, AXDND1, AXIN1, B3GNT1, B3GNT3, B4GALNT3, BAIAP2L2, BAMBI, BARX2, BCAS1, BCL10, BCMO1, BDKRB1, BLCAP, BMP2, BPNT1, BRI3, BSG, BTBD10, BTD, BTNL3, BTNL8, C10orf76, C11orf49, C11orf53, C11orf71, C11orf86, C12orf49, C15orf48, C19orf77, C1orf106, C1orf172, C1orf177, C1orf210, C21orf119, C2CD2L, C2orf72, C3orf52, C3orf83, C4orf19, C6orf141, C6orf47, C7orf43, C8G, C8orf82, C9orf69, CA13, CA4, CABLES1, CACFD1, CALM1, CALM2, CALML4, CAMSAP3, CAMTA1, CANT1, CAPN5, CAPN8, CASP16, CASP7, CASZ1, CBLC, CC2D1A, CCDC120, CCDC24, CCDC25, CCDC64B, CCDC85C, CCDC88C, CCDC96, CCL15, CCL28, CCNDBP1, CCNI2, CCNK, CCZ1B, CD151, CDC42, CDCP1, CDH1, CDH17, CDHR2, CDHR5, CDK16, CDKN1A, CDX1, CDYL, CEACAM5, CEP170B, CES2, CFDP1, CHMP1B, CHMP4B, CHMP4C, CHST6, CIB2, CKMT1A, CKMT1B, CLDN4, CLDN7, CLN3, CLRN3, CLSTN1, CMTM8, CNKSR1, CNOT11, CNPPD1, COASY, COL17A1, COMMD6, CORO2A, CRADD, CRAT, CRB3, CRK, CRYL1, CSNK1D, CSTF2, CTDSP1, CTNNA1, CTSA, CTSS, CTSV, CUX1, CYFIP1, CYP2C18, CYP2S1, CYP4F11, DAGLB, DAZAP2, DDAH2, DDR1, DDX19B, DECR2, DENND1A, DGAT1, DGCR2, DGKQ, DHCR24, DHDDS, DHRS1, DHRS9, DIRC2, DLG3, DLST, DMTN, DNAJA2, DNAJC5, DOK4, DOLPP1, DPP3, DUSP8, DYNC2LI1, ECI1, EFCAB4A, EFNA2, EFNA4, EFNB2, EGLN3, EIF2AK1, EIF4H, ELF3, ELMO3, ELOVL1, ELOVL6, EMP1, ENTPD2, ENTPD6, ENTPD8, EPCAM, EPPK1, EPS8L2, ERCC5, ERGIC1, ERN2, ESPN, ESRRA, ETHE1, EXD3, F11R, FAM102A, FAM118B, FAM132A, FAM189A2, FAM195A, FAM222B, FAM3A, FAM3B, FAM3D, FAM45A, FAM46A, FAM63A, FAM83E, FAM83F, FAM83G, FAXDC2, FBLIM1, FBXL12, FBXO31, FBXW4, FCGBP, FDFT1, FKBP1B, FLNB, FLVCR2, FOXA3, FRMD3, FTH1, FUT2, FUT3, FUT6, FXYD3, GABARAPL2, GAK, GAL3ST1, GALNT3, GALNT7, GBP3, GCNT3, GDPD2, GDPD3, GDPGP1, GFER, GGT6, GIPC1, GJB2, GLRA4, GNA11, GOLM1, GOLT1A, GPA33, GPER1, GPR108, GPR128, GPR153, GPR35, GPR39, GPRC5C, GPT, GSDMB, GSKIP, GSTO2, GTF3C5, GUCD1, HAAO, HAGH, HAS3, HDAC1, HDHD3, HEBP1, HEBP2, HECTD3, HEPH, HES1, HEXIM1, HIST1H1C, HIST1H2AE, HIST1H2AG, HIST1H2BD, HIST1H3D, HIST2H2BE, HIST2H4B, HKDC1, HMCN2, HMOX1, HNF4A, HOMEZ, HOOK2, HOXA11, HOXB8, HPS1, HRASLS2, HRCT1, HS1BP3, hsa-mir-1199, HSPA1B, HTATIP2, ICA1, IDNK, IFI27, IFI27L1, IFITM10, IFNGR2, IFT172, IGSF8, IHH, IL10RB, IL17RE, IL18, IL20RA, IL22RA1, IL32, INCA1, INF2, INPP5A, IQCK, IQSEC2, IRF6, ISCA2, IST1, ITFG1, ITGB6, ITLN1, ITM2C, ITPKA, ITPR3, JOSD1, JUP, KALRN, KAT5, KAZALD1, KCMF1, KCNK1, KCTD5, KDELR2, KIAA0319L, KIAA1161, KIAA1211L, KIAA1522, KLF4, KLF6, KLK15, L1TD1, LACTB2, LASP1, LDHD, LDLRAP1, LENG9, LGALS3, LGALS4, LGALS9, LHFPL2, LLGL2, LMAN2L, LMO7, LPAR5, LPCAT4, LPHN1, LRP5, LRRC16A, LRRC66, LSM14B, LSR, LUZP1, LXN, MACROD1, MAGIX, MALRD1, MANSC1, MAP1LC3A, MAPK13, MAPK3, MARK2, MAX, MBOAT7, MCF2L, MED18, MEP1A, MESP1, MFSD11, MFSD3, MGAT4B, MGAT5, MIEF2, MISP, MKNK2, MLX, MMAA, MMP15, MOGAT3, MPI, MPLKIP, MPND, MPST, MST1R, MTRF1L, MTUS1, MUC13, MUC20, MUC3A, MVK, MVP, MXD1, MYH14, MYL12A, MYL12B, MYO1C, MYO1E, MYO7B, NAA60, NADK, NAGPA, NANS, NAPRT1, NARFL, NAT2, NEDD9, NEK11, NFE2L1, NOSTRIN, NOXA1, NOXO1, NPAS2, NR2F6, NRL, NSMF, NT5C, NT5E, NTAN1, NUMB, OAS1, OASL, OGDH, OPLAH, OPTN, OSBPL2, OVOL1, OVOL2, PACSIN2, PAFAH2, PAK1, PAK4, PARD3, PARM1, PARP12, PAX8, PC, PCDH1, PCSK7, PCYT1A, PDGFA, PDLIM2, PERP, PET117, PEX7, PHGR1, PIGC, PIGR, PIGS, PINK1, PIP4K2C, PITPNM3, PLA2G10, PLA2G4F, PLAC8, PLCB3, PLCD3, PLEKHJ1, PLGRKT, PMEL, PNPLA2, POLD4, POLR2H, POLRMT, PON2, PON3, PPAP2C, PPARD, PPCS, PPIC, PPP1R14D, PPP1R16A, PPP1R26, PRAP1, PRKAG2, PRKCD, PRKCZ, PRR13, PRR15L, PRRG2, PRSS16, PRSS3, PRSS8, PSEN1, PSORS1C1, PTK6, PTPN18, PTPRH, PTPRN2, PTTG1IP, PYGB, RAB11A, RAB11FIP4, RAB19, RAB25, RAB3B, RAB40C, RAB9A, RABIF, RABL6, RAC1, RAC3, RALBP1, RASSF7, RBKS, RBM10, RCE1, RER1, RHBDL2, RHOV, RHPN2, RIMS3, RIPK1, RIPK3, RIPK4, RNF123, RNF185, RNF186, RNF223, RNF7, RPS6KA1, RUNDC1, RUSC1, S100A14, SAR1B, SBDS, SCAMP2, SCNN1A, SCRN2, SDC4, SDCBP2, SECTM1, SERAC1, SERINC2, SESN2, SETD3, SF3B4, SGPL1, SGPP2, SGSM3, SH2D3A, SH2D4A, SH3D21, SH3KBP1, SHD, SI, SIAE, SIGIRR, SIRT6, SIRT7, SKP1, SLC12A7, SLC15A1, SLC22A18, SLC22A18AS, SLC25A1, SLC25A5, SLC27A2, SLC29A4, SLC35A4, SLC35D2, SLC39A11, SLC39A4, SLC39A9, SLC44A2, SLC44A4, SLC45A3, SLC52A3, SLC5A9, SLC6A8, SLC7A9, SLC8B1, SLC9A1, SLC9A3R1, SMAD3, SMAP1, SMG7, SMIM22, SMIM6, SMPD1, SMPD3, SNX33, SORT1, SOX13, SP2, SPATA2, SPATS2L, SPDEF, SPECC1, SPHK2, SPINT1, SPINT2, SPRY2, SPTAN1, SPTBN1, SRC, SSBP3, SSH3, ST14, ST6GALNAC6, STAMBP, STAP2, STARD5, STAU1, STK24, STPG1, STXBP2, SUGT1, SYAP1, SYT7, SYTL5, TADA2B, TANGO2, TAX1BP1, TAX1BP3, TBC1D10A, TBC1D22B, TBCC, TBX10, TCTA, TEAD3, TFF3, TICAM1, TINAGL1, TJP2, TJP3, TLDC2, TM4SF5, TM6SF2, TM7SF2, TMBIM1, TMBIM6, TMC4, TMC7, TMCO1, TMCO4, TMED4, TMEM106C, TMEM125, TMEM127, TMEM139, TMEM14C, TMEM150B, TMEM159, TMEM164, TMEM180, TMEM184A, TMEM219, TMEM229B, TMEM242, TMEM246, TMEM253, TMEM45B, TMEM50A, TMEM51, TMEM53, TMEM54, TMEM61, TMEM63B, TMEM82, TMEM8A, TMPRSS2, TMPRSS4, TMX2, TNK1, TOB1, TOB2, TOE1, TOM1L2, TOR1B, TOR2A, TOX4, TP53I3, TPRG1L, TPRN, TPSG1, TRAF4, TRIM14, TRIM31, TRIP10, TRPT1, TSC22D1, TSEN54, TSKU, TSPAN1, TSPAN15, TSPAN3, TSPAN31, TSPAN8, TST, TTC22, TTC38, TTC39A, UCK1, UPF1, URGCP, USH1C, VAMP3, VAPA, VDAC1, VDAC2, VILL, VIPAS39, VPS26B, VPS28, VPS33B, VPS37B, VRK3, VSIG2, VWA3B, VWA7, WBSCR27, WDR55, WSB2, WWP2, XRCC4, YAE1D1, YWHAB, YWHAZ, ZBTB7A, ZBTB7B, ZDHHC3, ZDHHC5, ZER1, ZFAND2A, ZFAND2B, ZFPL1, ZFYVE1, ZFYVE21, ZFYVE28, ZMYND12, ZNF341, ZNF706, ZNF784, ZNF821, ZNRF1,* |
